# Generation of antigen-specific paired chain antibody sequences using large language models

**DOI:** 10.1101/2024.12.20.629482

**Authors:** Perry T. Wasdin, Nicole V. Johnson, Alexis K. Janke, Sofia Held, Toma M. Marinov, Gwen Jordaan, Léna Vandenabeele, Fani Pantouli, Rebecca A. Gillespie, Matthew J. Vukovich, Clinton M. Holt, Jeongryeol Kim, Grant Hansman, Jennifer Logue, Helen Y. Chu, Sarah F. Andrews, Masaru Kanekiyo, Giuseppe A. Sautto, Ted M. Ross, Daniel J. Sheward, Jason S. McLellan, Alexandra A. Abu-Shmais, Ivelin S. Georgiev

## Abstract

The traditional process of antibody discovery is limited by inefficiency, high costs, and low success rates. Recent approaches employing artificial intelligence (AI) have been developed to optimize existing antibodies and generate antibody sequences in a target-agnostic manner. In this work, we present MAGE (Monoclonal Antibody GEnerator), a sequence-based Protein Language Model (PLM) fine-tuned for the task of generating paired human variable heavy and light chain antibody sequences against targets of interest. We show that MAGE can generate novel and diverse antibody sequences with experimentally validated binding specificity against SARS-CoV-2, an emerging avian influenza H5N1, and respiratory syncytial virus A (RSV-A). MAGE represents a first-in-class model capable of designing human antibodies against multiple targets with no starting template.

## Main Text

Human monoclonal antibodies are a diverse class of therapeutics that can theoretically target any protein with exquisite specificity, making them promising candidates for treating a wide variety of diseases. Until recently, antibody development has been primarily driven by discovery-based experimental methods, typically through screening human or animal samples with prior exposure to an antigen target of interest. Even with recent developments that have drastically improved the throughput of antibody discovery methods, this process is laborious, slow, and cost-ineffective. The continued growth of the therapeutic market and range of applications for monoclonal antibodies presents an increased demand for *in silico* tools that accelerate and expand the capabilities of antibody discovery.

Recent breakthroughs in artificial intelligence (AI), most notably the unmatched performance of transformer-based Large Language Models (LLMs) and diffusion models on various tasks, have enabled a surge in computational approaches for antibody-related design tasks. Such methods include affinity maturation(*1, 2*), antibody redesign(*3–5*), and generation of single-domain antibodies(*6, 7*). However, no published methods have demonstrated the ability to design template-free, antigen-specific antibodies. Existing approaches are limited to antibody redesign, with a focus on generation of complementarity-determining regions (CDRs), requiring an initial antibody template to provide variable genes and framework regions for the antibody. Additionally, such models are primarily structure-based and require antibody-antigen complexes for training, which is significantly limiting due to insufficient data, especially in the context of paired, human antibodies.

In this manuscript, we present MAGE (Monoclonal Antibody GEnerator), a protein language model capable of generating paired heavy and light chain antibody variable sequences with binding specificity against input antigen sequences. MAGE was developed by finetuning an auto-regressive decoder LLM that was pretrained on general protein sequences. Such models learn from observed amino acid sequences by next-token prediction, using self-attention to capture complex dependencies within input sequences. Here, we leveraged this learned representation of amino acid sequences as a starting point for learning human antibody sequence features associated with binding specificity to diverse antigen targets. We show that MAGE is capable of generating antibodies that exhibit diverse sequence features, including heavy and light chain variable gene usage, levels of somatic hypermutation (SHM), and novel CDRs not observed in the training data. When prompted with SARS-CoV-2 wildtype receptor binding domain (RBD), binding specificity was successfully confirmed for 9/20 of experimentally validated MAGE-generated antibodies, including one antibody with better than 10 ng/mL potency of SARS-CoV-2 neutralization. Binding antibodies were also designed and validated against RSV-A prefusion F (7/23 antibodies), which was significantly less represented in the training data. We determined a cryo-EM structure of two MAGE-designed antibodies in complex with RSV F, demonstrating that MAGE generates antibodies with diverse binding modes and can incorporate impactful residues at key binding interfaces. Finally, MAGE-designed antibodies were validated against H5/TX/24 hemagglutinin (HA) (5/18 antibodies), demonstrating zero-shot learning capabilities against an influenza virus strain that was not present in the training data. MAGE therefore represents a first-in-class model capable of designing novel human antibodies with demonstrated functionality against antigen targets of interest, without having to provide any part of the antibody sequence as a starting template.

### Fine-tuning a PLM for antigen-specific antibody generation

Here, we present a protein language model (PLM) called MAGE, fine-tuned for generating paired heavy and light chain antibody variable sequences that bind to a prompted antigen sequence. Toward this goal, a pretrained model was finetuned on a training database containing antibody-antigen sequence pairs curated from literature and existing databases (**Fig. 1a**). In addition to published data, we collected an original dataset of antigen-specific antibody sequences against diverse viral antigens using LIBRA-seq (Linking B-cell Receptors to Antigen-specificity through Sequencing), a high-throughput method for identification of antigen-specific B cell receptors (BCRs) against an antigen panel(*13*). A panel of 18 diverse antigens was used to screen peripheral blood mononuclear cells (PBMCs) from 20 donors distributed across four groups (HIV infected, influenza vaccinated, COVID-19 convalescent, and healthy). Using this diverse training dataset, we aimed to present a model capable of generating functional, target-specific antibodies against input antigen sequences. General protein language models have been shown to have superior performance on antibody-specific tasks due to the complex nature of understanding the input antigens, antibodies, and the interactions between them (*11*). We therefore finetuned a general protein model for the task of antigen-specific antibody generation.

**Figure 1.**
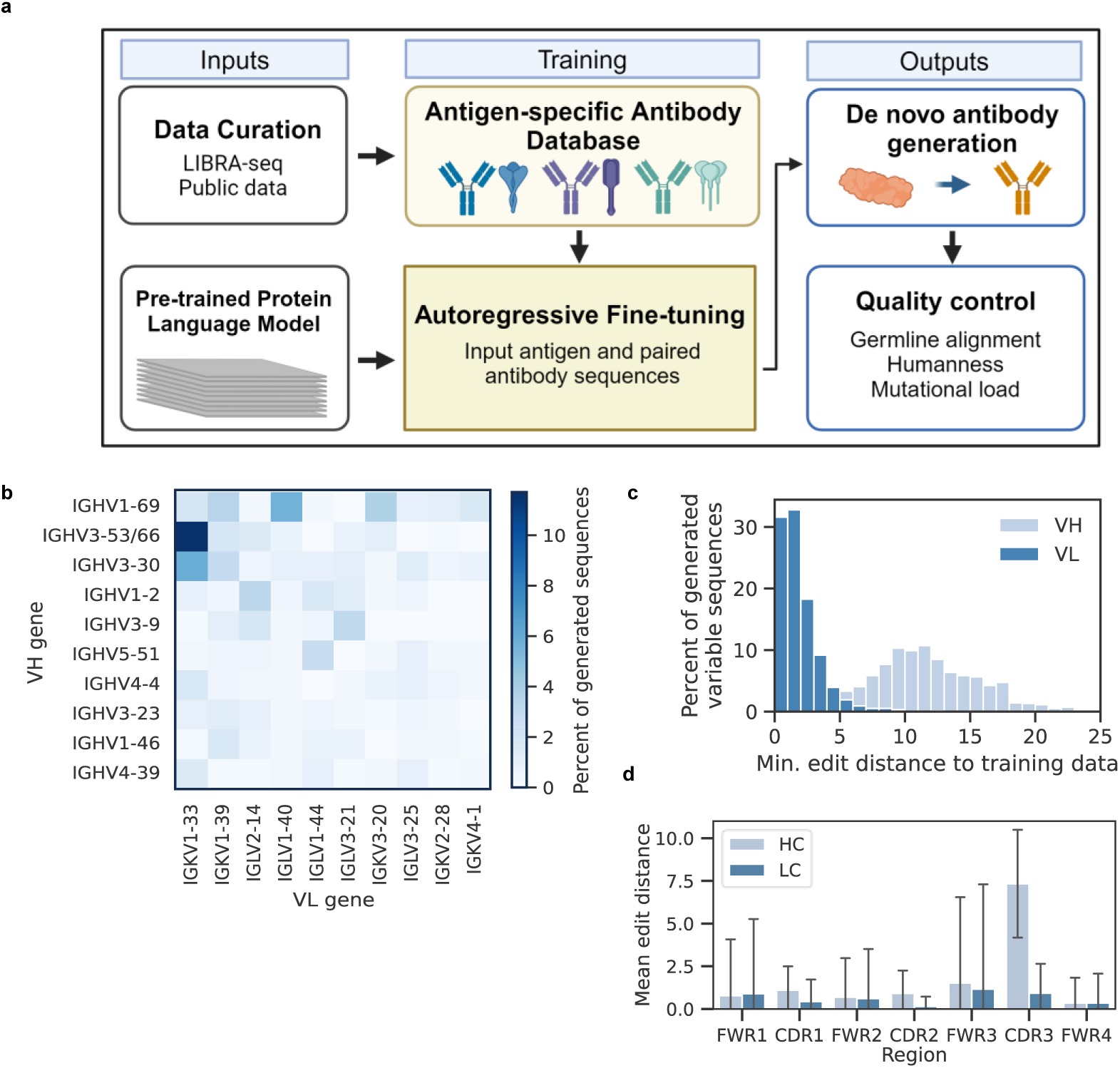
A general PLM was fine-tuned for antigen-specific antibody generation. A) An Antigen-specific Antibody Database was curated, in combination with large scale LIBRA-seq datasets, in order to fine-tune the pretrained PLM for paired chain antibody generation against antigen prompts. B) Percentage of 1,000 antibodies generated against RBD that use each combination of heavy and light V genes. The top 10 most common genes are shown for each. C) Generated variable heavy (VH) and variable light (VL) sequences were aligned to the training data to find the minimum number of mutations between each generated sequence and any training sequence. D) For the most similar training sequence from the comparison in C, the distance was calculated between each region of the VH or VL sequence. The mean across all RBD generated sequences are shown with error bars representing the standard deviation.

### Generated antibody sequences are diverse and distinct from training data

Following finetuning, the trained model could be prompted with an antigen sequence of interest to generate an output containing an antibody variable heavy and light chain sequence. To evaluate the ability of MAGE to generate antigen-specific antibody sequences, we selected three targets that spanned the range of training data representation. We first tested generation against SARS-CoV-2 RBD, which had disproportionately higher representation in the training data. Then, to show that the model can successfully work for antigen targets with less training data, we tested against two additional antigens. To assess the quality and diversity of sequences generated by MAGE, 1,000 antibody sequences were generated against RBD and aligned to a human germline reference using IMGT numbering(*25*) and then filtered using the following criteria (described in detail in Methods): 1) Removal of sequences without a recognizable heavy or light chain, 2) removal of sequences with any missing CDRs or framework regions (FWRs), and 3) removal of variable heavy or light sequences less than 100 amino acids in length. Almost all (991/1,000) of the generated sequences passed these filters. Additionally, sequences were scored for ‘humanness’ using the open-source platform BioPhi OASis(*26*). Based on suggested thresholds, sequences with an OASis percentile score less than 70% were removed, with only 2.2% (22/991) sequences falling below this humanness threshold (**Fig. S2a)**. While these sequences could represent viable, particularly novel sequences, this model was intended to generate human antibodies for further characterization and these low-scoring antibodies by OASis were removed accordingly. In total, 969 of 1,000 generated sequences were retained for further analysis and down-selection for *in vitro* characterization.

The RBD-prompted sequences displayed diverse sequence features, using 37 unique variable heavy chain genes and 30 unique variable light chain genes, not accounting for different alleles. In total, 322 different pairs of heavy and light variable genes were represented in the generated sequences, with the most frequently used pair (IGHV3-53/66: IGKV1-33) representing only 13.9% (135/969) of sequences (**Fig. 1b**). Generated sequences also showed diverse CDRs, with heavy chain CDR3 (CDRH3) lengths ranging from 5 to 28 amino acids (mean = 16), and light chain CDR3s (CDRL3) lengths ranging from 7 to 12 amino acids (mean = 10) (**Fig. S2b**). The light chains were more biased towards germline, with 50.1% (486/969) of containing no mutations, compared to 18.1% (175/969) for the heavy chains (**Fig. S2c**). These results suggest that rather than simply using a single dominant heavy-light chain combination, MAGE is capable of generating diverse populations of antibody sequences.

We next sought to determine the novelty of generated antibodies at an individual level. In an attempt to quantify this novelty, each generated sequence was compared to all sequences seen during fine-tuning to identify the most similar training example based on the minimum Levenshtein distance between each pair of sequences. This distance, which can be intuitively interpreted as the number of amino acid differences, was first calculated separately for the heavy and light chains (**Fig. 1c**). We observed that the generated heavy chains contained more differences from training data sequences on average (mean = 11.7 differences), compared to light chains which exhibited substantially lower levels of differences (mean = 1.4 differences). When separated based on antibody sequence region, the distances to the nearest training sequence were highest in the CDR3s (**Fig. 1d)**, as could be expected due to the high diversity in this region. Distances were higher for the heavy chains than light chains across all regions aside from framework region 4 (FWR4). We also compared the similarity of generated heavy chain CDRH3s specifically to the training RBD sequences by finding the maximum sequence identity based on Levenshtein distance. The generated CDRH3s were largely novel, covering a range of identities centered at a mean of 72.4% sequence identity, with 7.4% of the generated antibodies containing CDRH3s identical to an antibody seen in training (**Fig. S2e**). The distribution of similarity to training data based on both heavy and light chain distance and CDR3 identity was broad **Fig. S2d**), suggesting that the generated antibody sequences cover a range of uniqueness with respect to sequences seen in training. Together, these results indicated that MAGE generated sequences with differences in all regions of the antibody, rather than only designing CDRs.

### Generated antibodies exhibit diverse binding profiles to SARS-CoV-2 RBD

Following basic filtering of the 1,000 generated antibody sequences, we used a simple pipeline to select antibodies for experimental validation of binding (**Fig. 2a**). From the 969 antibody sequences that remained after filtering, 10 antibodies were first chosen in an unbiased manner, without comparison against RBD-specific antibodies: to test a diverse unbiased set, the 20^th^ and 80^th^ percentile of VH germline identity antibodies from sequences using the top 5 most frequently generated VH genes were selected. Another set of 10 antibodies was selected based on similarity to known RBD-specific antibodies: for this biased selection, the top 5 antibodies with the highest CDRH3 identity to any CoV-AbDab antibody, and top 5 antibodies with the highest VH identity to any CoV-AbDab antibody were chosen. In total, a set of 20 antibodies was selected for *in vitro* validation, containing a range of sequence characteristics and novelty which aimed to represent the distribution of generated sequences (**Supplemental Table 2**). When compared to the most similar training antibody, the selected antibodies ranged from a minimum VH distance of 3 residues (RBD-238) to 24 different residues (RBD-153) (**Fig. 2b)**. The respective light chains were more similar to those seen in training, with a minimum distance ranging from 0 residues (RBD-135) to 9 residues (RBD-727).

**Figure 2.**
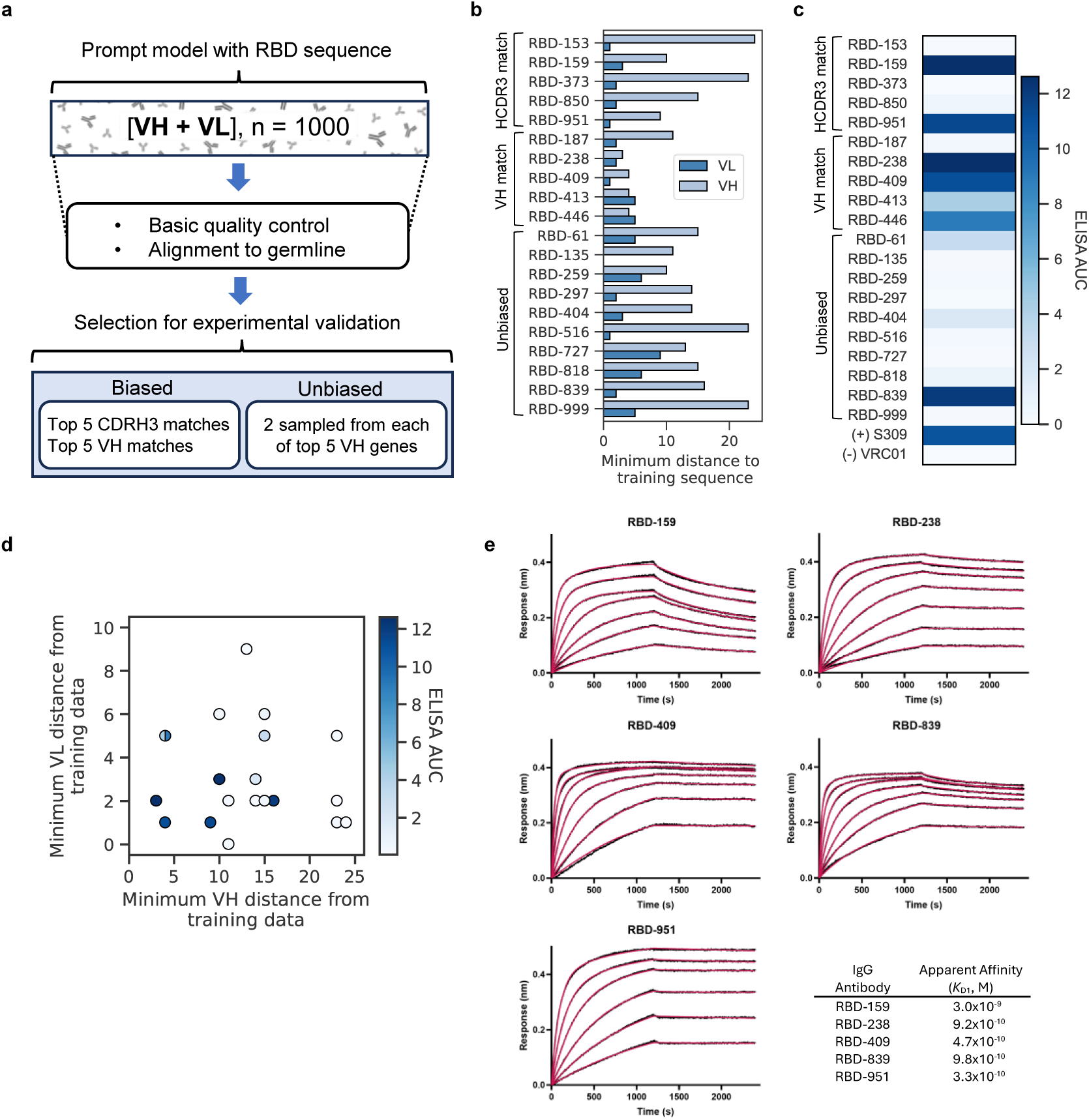
Twenty antibodies were selected for experimental validation of binding to RBD. A) Schematic of antibody selection method after generation, yielding a total of 20 antibodies for experimental validation. B) For each antibody, the Levenshtein distance for the VH or VL is shown in comparison to the training antibody with the lowest total distance (summed across VH and VL). Antibodies are grouped by selection group. C) ELISA area-under-the-curve (AUC) based on absorbance at 450 nm across a dilution series from 6.4×10-4 μg/mL to 10 μg/mL, with S309 (RBD-specific) positive control and VRC01 (HIV-1-specific) negative control antibodies. D) Relationship between the minimum VH and VLs distance from the closest training antibody sequences with points colored based on ELISA AUC. Overlapping points at VH distance = 4 and VL distance 5 are shown a single point with split coloring based on AUCs of these two antibodies (RBD-446, RBD-413). E) BLI sensorgrams for binding of high-affinity IgG antibodies to immobilized SARS-CoV-2 RBD-SD1. Data (black) were fit to a 1:2 bivalent analyte model. Curve fits are shown in red.

The 20 antibodies selected for experimental validation were tested for binding to RBD from the SARS-CoV-2 index strain by ELISA (enzyme-linked immunosorbent assay) (**Fig. 2c, Fig. S3**). From these results, 9/20 (45%) of the tested antibodies were identified as binding hits for further characterization based on a minimum of 2-fold signal over background at the highest antibody concentration tested (10 μg/mL). In the biased selection group, 2/5 of the CDRH3 matches (RBD-159, RBD-951) and 4/5 of the VH matches (RBD-238, RBD-409, RBD-413, RBD-446) displayed binding by ELISA. In the unbiased selection group, 3/10 antibodies (RBD-61, RBD-404, RBD-839) displayed binding, with RBD-839 displaying the strong binding signal, on par with the positive control antibody S309(*27*). All of these binding antibodies showed no detectable ELISA signal to BG505, an HIV-1 envelope trimer. While the binding antibodies generally exhibited lower minimum distances from both VH and VL training sequences compared to the antibodies that showed no binding, the binding antibodies nevertheless exhibited substantial novelty, with a range of 5-25 (mean: 13.6) total distance to closest training antibody (**Fig. 2d**). In particular, RBD-839 showed a higher minimum distance from the nearest training antibody (total distance = 18 residues) than 67% of the non-binding antibodies (**Fig. 2d**). We observed a wider range of VH distances to training data compared to VL, in alignment with the lower diversity of light chains we previously observed in the pool of generated antibodies and training data.

Binding was further validated using biolayer interferometry (BLI) to measure association and dissociation kinetics for IgG binding to immobilized, monomeric SARS-CoV-2 RBD (RBD-SD1) (**Fig. 2e, Fig. S4)**. Apparent *K*_D_ (*K*_D1_) values were determined by fitting the resulting binding curves to a 1:2 bivalent analyte model(*28*), which accounts for the slower observed dissociation rate due to avidity. Of the hits identified by ELISA, 8 of 9 demonstrated measurable binding to RBD-SD1, with no binding observed for RBD-404 at the highest concentration tested (1,024 nM). Four antibodies from the biased-selection groups (RBD-159, RBD-238, RBD-409, and RBD-951) and one from the unbiased-selection group (RBD-839) demonstrated apparent high-affinity binding, with *K*_D1_ values in the nanomolar to sub-nanomolar range.

RBD-61, RBD-413, and RBD-446 also bound to RBD-SD1, albeit with reduced apparent affinity (**Fig. S5**). Although a small amount of non-specific binding was detected for one antibody (RBD-951, **Fig. S4**), for the other 7/8 antibodies no binding was observed by BLI to a prefusion-stabilized RSV F trimer (DS-Cav1(*29*)), which is consistent with the specificity observed by ELISA.

Although Figure 2d shows that the antibody sequences are distinct from the training data, exhibiting a range of novelty, we sought to assess how similar the generated binding antibodies are at the population level. Public antibody clones, commonly defined by matching variable genes and CDR3 identity >70%, represent a set of criteria for grouping similar antibodies found in different individuals that are likely to share the same binding specificity(*30–32*). When comparing the generated binding antibodies to antibodies from the CoV-AbDab using this definition, we found a range of ‘publicness’, from zero public clones for RBD-61 and RBD-404 to the highly public RBD-238 with over 100 public clones (**Fig. 3a**). We observed that all generated binding antibodies had >70% CDRH3 identity with at least one CoV-AbDab antibody, which is not surprising given the vast diversity and size of the database. Some of these antibodies, e.g. RBD-951, shared sequence features with many training antibodies at a population level, while others, e.g. RBD-61, appeared much less public (**Fig. S5**). When comparing each generated binding antibody to its closest training match based on VH distance, we observed that the majority of differences were in the CDRH3, but almost all (8/9) of the binding antibodies contained at least one difference outside of the CDRH3 (**Fig. 3d**), demonstrating the ability of MAGE to generate distinct full variable sequences rather than only designing CDRs. In addition to varying levels of publicness and locations of mutations, we demonstrate that RBD-specific antibodies generated by MAGE have diverse sequence features including CDR lengths, variable gene usage, and humanness scores (**Fig. 3c**).

**Figure 3.**
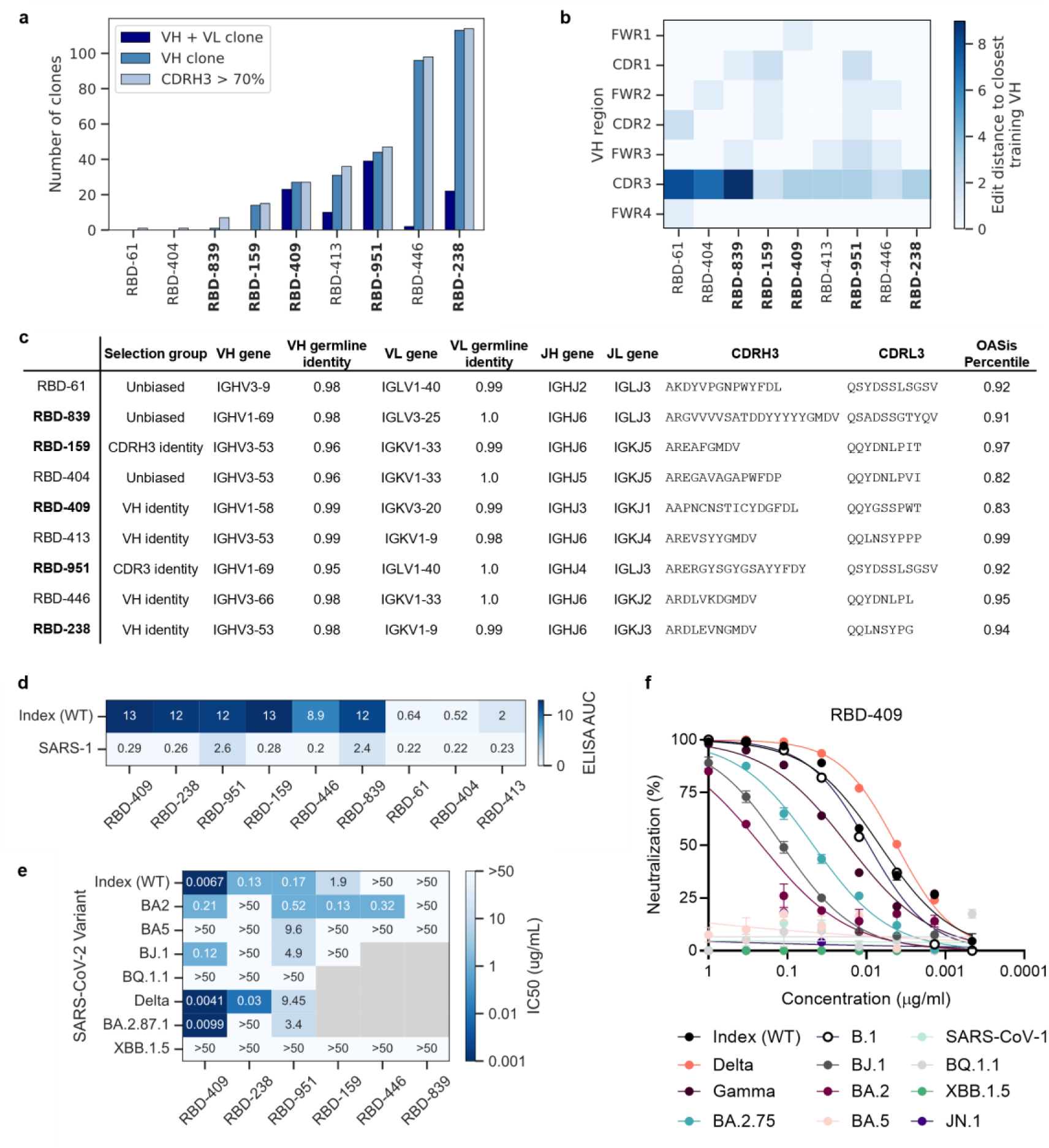
Generated RBD binding antibodies have diverse sequence characteristics. Strong affinity binding antibodies are bolded. A) Publicness of binding antibodies based on CDRH3, VH, and both VH and VL. Clones defined as CDR3 identity > 70% and matching V genes for the specified chain. B) Sum of edits within each VH region for binding antibodies compared to closest sequence match in training data. C) Table showing sequence characteristics of RBD binding antibodies. OASis percentile represents a humanness score averaged across the heavy and light chains D) ELISA AUC for binding curve dilutions for SARS-CoV-2 WT and SARS-CoV-2 spikes for RBD binding antibodies. E) IC50 values for psuedovirus neutralization of SARS-CoV-2 variants for full spike binding antibodies. F) Full pseudovirus neutralization curves for RBD-409 against SARS-CoV-2 variants.

### Generated RBD antibodies bind full-length spike and neutralize SARS-CoV-2

The 9 binding antibodies to RBD were tested for binding to full-length SARS-CoV2 spike (index), along with a highly mutated variant (XBB.1) and SARS-CoV spike (SARS-1). Although MAGE was prompted using RBD only, we wanted to interrogate whether the generated antibodies would be compatible with and bind full-length spike. Of the RBD binding antibodies, 3/9 showed low to no binding to full-length spike in ELISA, suggesting that these antibodies may bind epitopes that are occluded or bind in conformations that may be sterically hindered on the spike trimers (**Fig. 3d**). All of the 6/9 antibodies which did bind to full-length spike also displayed binding to XBB.1, and two also displayed weak signal to SARS-CoV spike. These results further emphasize that MAGE was able to generate antibodies with diverse characteristics and binding properties, exhibiting cross-reactivity to different coronavirus spike variants.

Following validation of binding by ELISA, we aimed to determine whether the generated antibodies also exhibited virus neutralization in a pseudovirus assay(*32*). Four of the RBD binding antibodies displayed neutralization against SARS-CoV-2 index pseudovirus, with RBD-409 displaying highly potent neutralization (IC_50_ = 6.7ng/mL) (**Fig. 3e, Fig. S6**). Out of the 6 antibodies that bound full spike in ELISA, all but one showed neutralization potency of <1μg/mL against at least one spike variant. None of these antibodies were able to neutralize XBB.1.5, although this was unseen in training as the newest RBD variant included in training was Omicron BA.5. Nevertheless, RBD-409 displayed high neutralization potency against the SARS-CoV-2 spike Gamma (IC_50_ = 17ng/mL) and Delta (IC_50_ = 4.1 ng/mL) variants and was able to retain neutralization against several Omicron variants including BA.2, BA.2.75, and BJ.1 (**Fig. 3f**).

### MAGE is capable of generating functional antibodies against diverse targets with lower representation in the training datasets

While the training dataset used to fine-tune MAGE was highly biased towards coronavirus antibody-antigen pairs, the dataset did contain other diverse antigen specificities to enable generation against different prompts. To that end, antibodies were designed and tested for binding against a newly emerging highly pathogenic avian influenza virus (*33*) (H5) and the RSV-A glycoprotein prefusion F (RSV-A). For RSV-A, there were 292 training antibodies against the exact RSV F sequence used for prompting, along with 753 antibodies against related RSV antigens including RSV-B and post-fusion RSV F. Hence, the number of exact prompt training antibodies for RSV-A represented approximately 1/10 of the size of the training antibodies for SARS-CoV-2 RBD. In addition to validating antibody designs against a target with limited training data, we also sought to test the capability of MAGE to generate antibodies against a target not seen in training (zero-shot). Toward this goal, we prompted MAGE using hemagglutinin from the avian influenza (H5N1 clade 2.3.4.4b virus), an emerging public health threat with multiple reported interspecies transmissions, including human infections(*33, 34*). While this exact antigen sequence was not seen in training and was not even reported at the time of training MAGE, a total of 472 H5N1-specific antibodies were included in training. These training antibodies were primarily specific to the hemagglutinin variant A/Indonesia/05/2005(*17*), which has 91.5% sequence identity to the more recent A/Texas/37/2024 used for prompting. This target therefore represents a realistic use-case where MAGE can generate antibodies against an emerging threat without pre-existing knowledge of binding antibodies for that specific target antigen sequence.

To explore the behavior of MAGE when prompted with different antigens, 1,000 sequences were generated against A/cattle/TX/2024 H5 and RSV-A F. Notably, there was a significant enrichment of RSV-A and H5-specific clones generated when prompting with the respective antigens as opposed to prompting with SARS-CoV-2 RBD (**Fig. 4a**), suggesting that MAGE can enrich for prompt-specific antibody sequences. Further, each of the three prompts yielded antibody sequences with unique distributions of VH gene usage, with antibodies generated with the SARS-CoV-2 RBD prompt most frequently using IGHV3-53/66, in alignment with reported gene usage biases in SARS-CoV-2-specific repertoires(*35*), while RSV-A sequences heavily biased towards IGHV1-18 and H5 sequences towards IGHV4-34 (**Fig. 4b**). The antibody sequences generated against each prompt were then compared to the training data to find the minimum Levenshtein distance for each heavy and light chain, indicating that H5 and RSV-A prompted antibodies were more novel, on average, than the RBD prompted antibodies (**Fig. 4c-d**). Additionally, we found that the H5 and RSV-A prompted sequences exhibited higher levels of somatic hypermutation (SHM) than the RBD-prompted antibodies (**Fig. 4e-f**).

**Figure 4.**
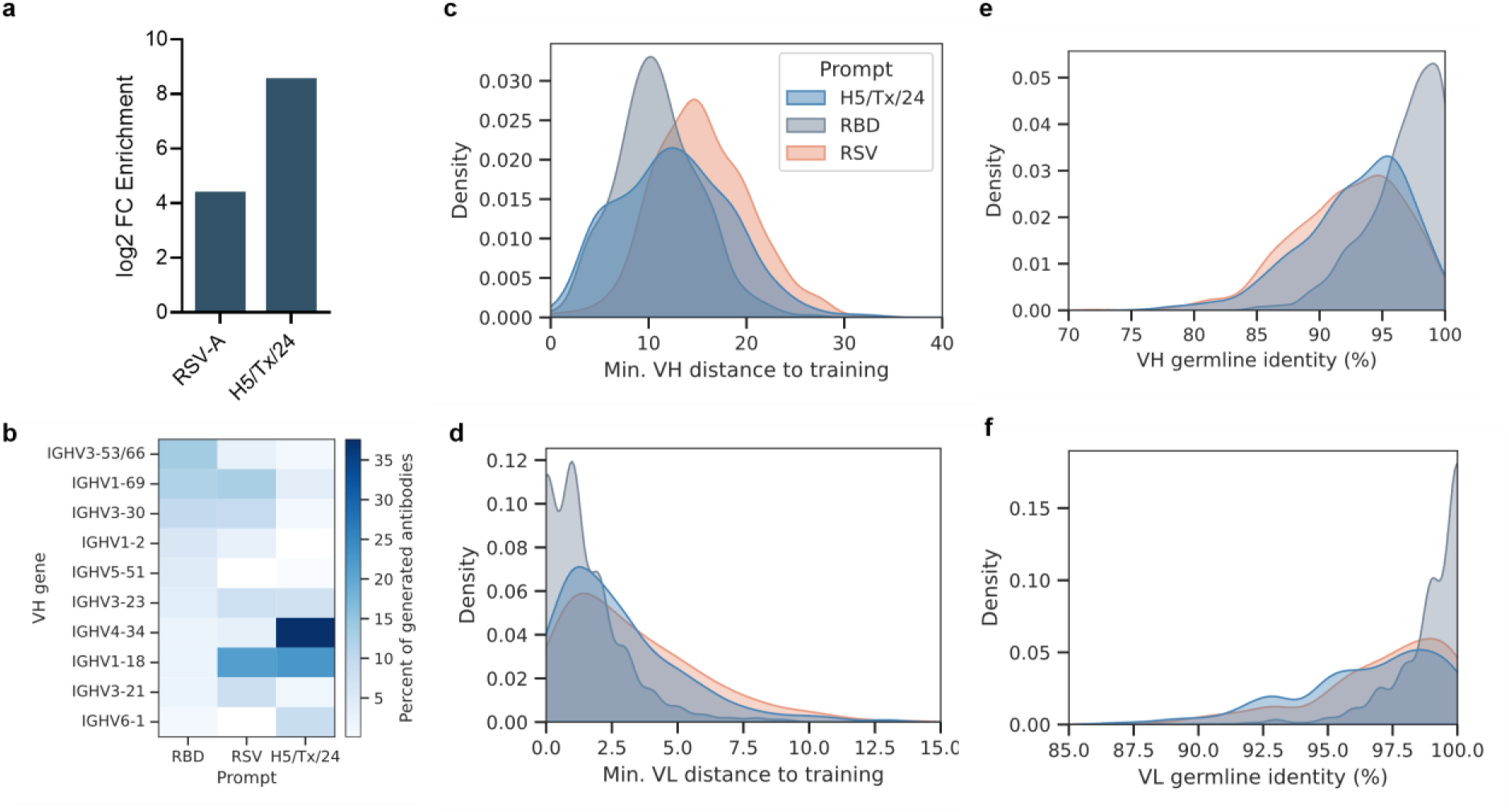
Characteristics of sequences generated against RSV and H5/TX/24 prompts. A) Log fold changes showing increase in same antigen-specificity clones for RSV-A and H5/TX/24 prompts compared to WT RBD prompt. Calculated based on number of clones between generated and training antibodies, out of 1000 generated sequences. B) Heatmap showing percent of 1000 generated antibody encoding different variable genes for each antigen prompt. For 1000 generated antibodies against each prompt, the distribution of C) minimum VH Levenshtein distance to any training antibody, D) minimum VL Levenshtein distance to any training antibody, E) percent identity to VH germline, and F) percent identity to VL germline.

We next sought to experimentally validate the binding specificity for a subset of these generated sequences against the H5 and RSV-A prompts. For H5, we compared the generated sequences to H5 training antibodies and selected a validation set of 18 designed sequences for experimental validation, aiming to capture a range of novelty compared to the training examples seen (see methods section - *Antibody selection for experimental validation of H5N1 antibodies,* **Table S2**). We confirmed strong binding by ELISA for 5/18 (28%) of these designs (**Fig. 5a**), along with another seven weak binding antibodies (>2-fold signal over background and >0.5 absorbance) at 10 µg/mL ELISA (**Fig. S7a**). The minimum distance to training antibodies for the binding antibodies ranged from 4-16 residues for the heavy chain, and 1-8 residues for the light chain (**Fig. 5b**). The levels of SHM ranged from 6-11% for the heavy chain, and 7-8% for the light chain (**Fig. 5c**). Similarly to the antibodies designed against RBD, novel residues in these H5-prompted antibodies were found across the entire VH region (**Fig. 5d**) and were not limited to the CDRs. Notably, all five of the strong H5 binding antibodies were neutralizing against influenza strains A/Texas/37/2024, A/Vietnam/1203/2004, and A/Michigan/45/2015 (**Fig. 5e, 5f, Fig. S7e-f**). Antibodies H5-242 and H5-384 were the most potent (IC_50_ < 100 ng/mL, **Fig. 5f**), with IC_50_s comparable to the positive control CR9114, a potently and broadly neutralizing stem-binding antibody(*36*).

**Figure 5.**
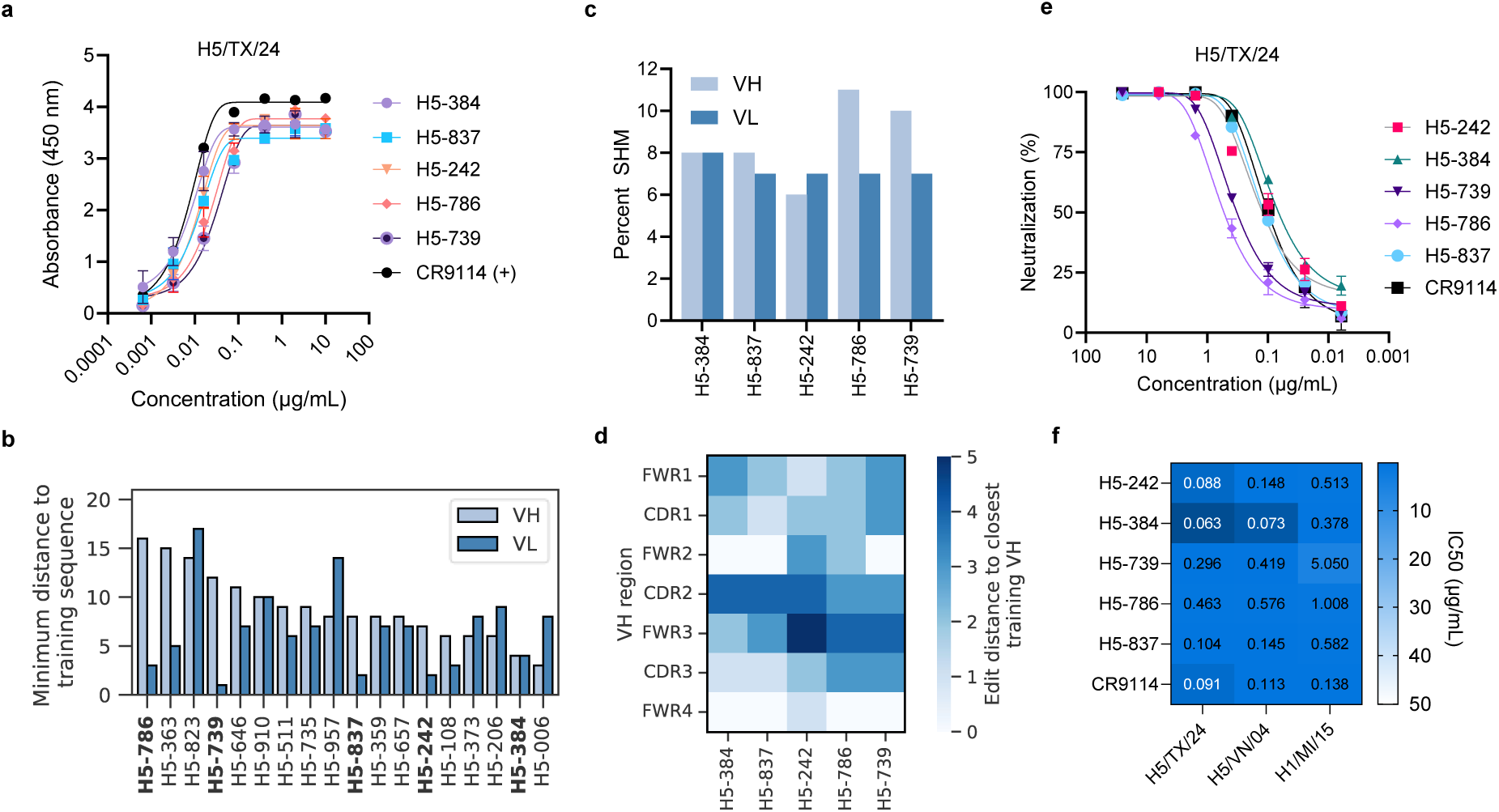
MAGE generates novel A/cattle/TX/2024 H5-binding antibodies. A) Full ELISA dilution curves for designed antibodies against H5/TX/24 hemagglutinin. B) Minimum distance to training antibody sequences. Distance represents number of residues different when compared to the heavy and light chain sequences from the training match with the lowest total distance (VH + VL). C) Percent somatic hypermutation in heavy and light chain for binding antibodies, calculated across VH and VL genes. D) Edit distance by VH region to closest training sequence match based on CDRH3 identity. E) Neutralization dilution curves against H5/TX/24 hemagglutinin. F) IC_50_ values calculated from curves shown in panel E.

For the RSV-A prompt, we generated a larger pool of 10,000 antibodies, followed by selection for validation of biased and unbiased selections using a similar stratification method as used for RBD (see methods section - *Antibody selection for experimental validation of RSV antibodies*), yielding a set of 23 antibodies for experimental validation of binding (**Table S2**). Following initial screening (**Fig. S7c**), we confirmed binding by ELISA for 7/23 (30%) of these designs, including three antibodies that were selected without biasing towards known RSV-specific antibodies (**Fig. 6a**). While the seven binding antibodies had at least one heavy chain clone in the training data (>70% CDRH3 identity, same VH gene), they nevertheless included many variations throughout the heavy and light chains ranging from a minimum heavy chain distance of six residues for RSV-6479 to 21 residues for RSV-2954 (**Fig. 6b**). In the light chain, the distances compared to training sequences range from 4 for RSV-4314 to 12 for RSV-3301. The MAGE-designed RSV binding antibodies showed SHM levels ranging from 3-21% for the heavy chain, and 2-12% for the light chain variable region (**Fig. 6c**), suggesting that MAGE does not simply learn germline-level antibody sequences. Compared to the closest training antibodies, we see that the RSV binding antibodies included a range of differences across the VH regions, including differences in at least 4/7 regions for all seven binding antibodies (**Fig. 6d**). There was also a range of novelty for the light chains in this set of antibodies, with the minimum VL distance to training antibodies ranging from 4-12 residues. The binding antibodies were further characterized by pseudovirus neutralization assays (**Fig. 6e**). Notably, 3/7 of the binding antibodies were able to neutralize RSV-A (**Fig. 6e**); while IC_50_ values were not determined, RSV-2245 and RSV-4314 appeared to be strongly neutralizing with neutralization >50% at 0.1 µg/mL. Notably, RSV-2245 was from the unbiased selection group, with a VH distance of 17 amino acids to the closest training antibody and a SHM level of 10%, representing a highly mutated antibody with a notably distinct sequence.

**Figure 6.**
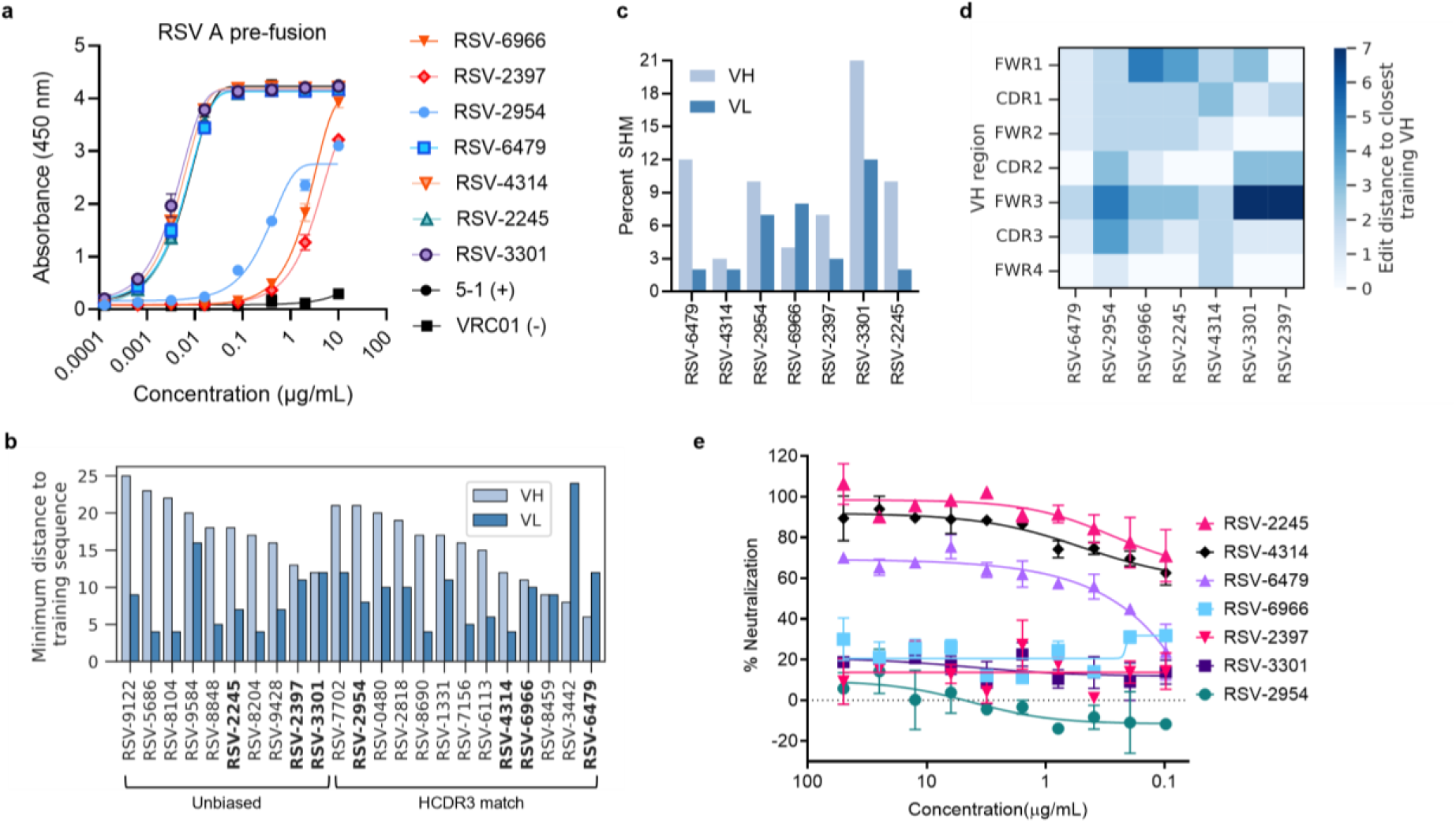
MAGE generates novel RSV-A binding antibodies. A) Full ELISA dilution curves for designed antibodies against RSV-A pre-fusion. B) Minimum distance to training antibody sequences. Distance represents number of residues different when compared to the heavy and light chain sequences from the training match with the lowest total distance (VH + VL). C) Percent somatic hypermutation in heavy and light chain for binding antibodies, calculated across VH and VL genes. D) Distance by VH region to closest training sequence match from E) Antibody neutralization dilution curves against RSV-A.

To investigate the epitopes targeted by MAGE-generated antibodies from the unbiased selection group with both high levels of SHM and high distances from training, we determined a cryo-EM structure of RSV prefusion F (PR-DM(*37*)) bound to fragments of antigen binding (Fabs) for RSV-2245 and RSV-3301 (**Fig. 7a**). For this complex, 140,634 particles were extracted from 1,323 micrographs to generate a 3.4 Å resolution reconstruction with 3 copies of each Fab bound to the RSV F trimer. The structure revealed that RSV-2245 binds to an epitope primarily within prefusion-specific antigenic site V, burying 850 Å^2^ on a single F protomer. Antibodies that target Site V are common in the human repertoire and tend to be potently neutralizing(*12*), consistent with the neutralization efficacy we observed for RSV-2245. RSV preF is contacted by all three CDRs of the RSV-2245 heavy chain and CDRs 1 and 2 of the light chain. The interface is centered on the strands of the β3-β4 hairpin, with a large network of hydrophobic contacts mediated by CDRH3 and Tyr53 of CDRH2. The sidechain of Tyr53_CDRH2_ additionally contacts a single residue within β2, forming a hydrogen bond with the sidechain of Tyr53_F_. The RSV-2245 light chain contributes additional interactions within β4 and with residues that flank the β3-β4 hairpin. Of note, Asp30_CDRL1_, which is mutated from asparagine in the germline sequence, forms a salt bridge with the Lys192_F_ sidechain. This mutation was only observed in 1/292 (0.34%) of the training RSV-specific antibodies, with the corresponding training antibody showing low similarity (73% LCDR1 identity and only 50% CDRH3 identity), demonstrating the ability of the model to learn sequence features from individual training sequences and integrate them into novel antibodies. The RSV-2245 epitope further extends to include residues within antigenic Site II, mediated by polar contacts between CDRH2 and the helix-turn-helix formed by the α6 and α7 helices.

**Figure 7.**
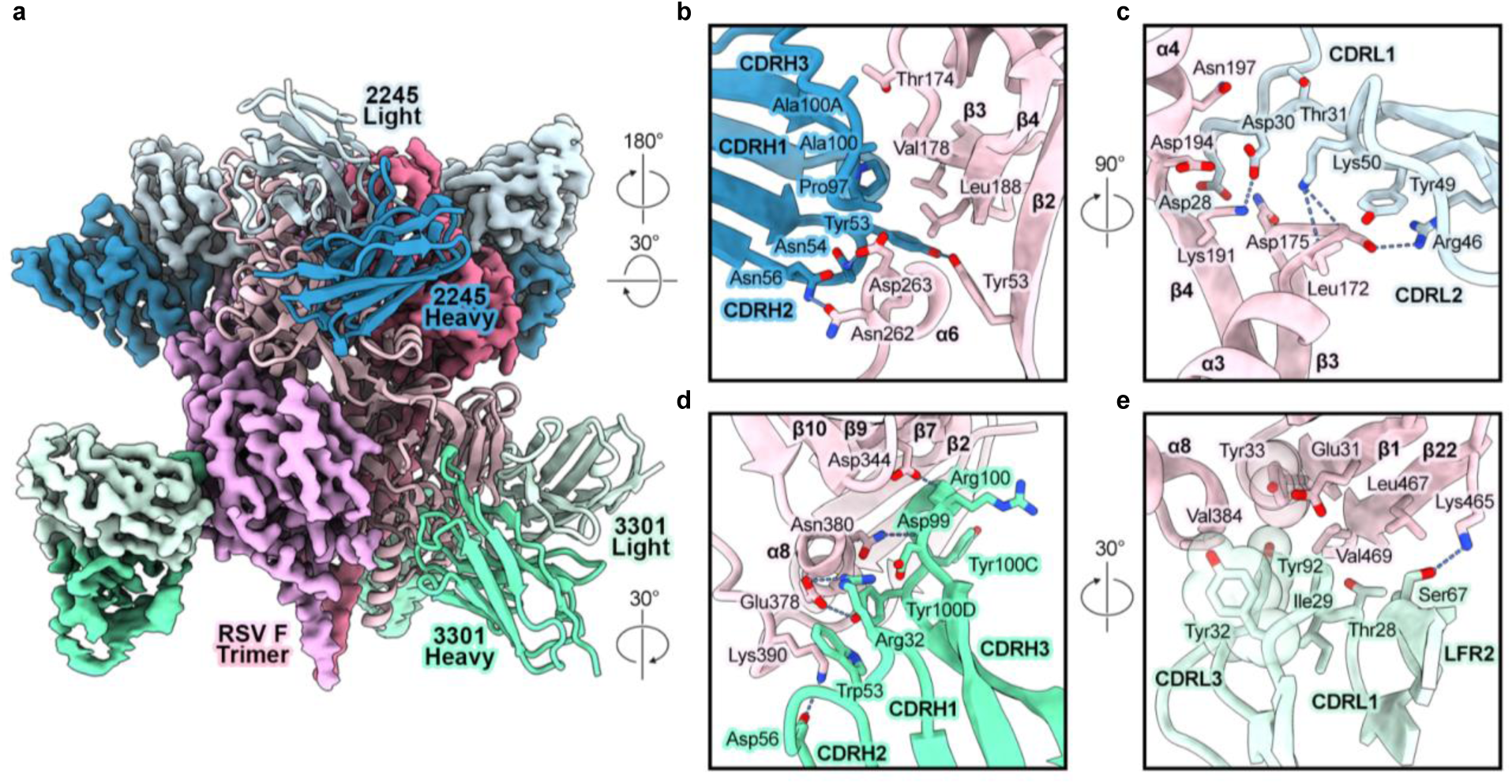
Cryo-EM structure of Fabs RSV-2245 and RSV-3301 bound to RSV-A. **F.** A) Overview of 3.4 Å resolution cryo-EM structure of RSV F bound to fragments of antigen binding (Fabs) for RSV-2245 (heavy and light chains in dark and light blue, respectively) and RSV-3301 (heavy and light chains in dark and light green, respectively). RSV-A F protomers are shown in shades of pink. Zoomed-in views of the Fab-RSV F interface are shown as cartoons with select residues represented as sticks for B) RSV-2245 heavy chain, C) RSV-2245 light chain, D) RSV-3301 heavy chain, and E) RSV-3301 light chain. Hydrogen bonds are shown as dashed blue lines.

RSV-3301 represents the most highly mutated antibody of the validated RSV-specific set. The structure revealed that RSV-3301 buries approximately 715 Å^2^ within the membrane-proximal lobe of one F protomer, targeting an epitope that lies almost entirely within antigenic Site I. This site is typically considered to be postfusion F-specific but is largely conserved in both pre- and postfusion conformations(*38–40*). The interaction is dominated by CDRH3, which extends into the cavity formed between the α8 helix and the curved β-sheet formed in part by β10, β9, β7, and β2. Backbone atoms within Asp99_CDRH3_ and Arg100_CDRH3_ form hydrogen bonds with the sidechains of Asn380_F_ and Asp344_F_, respectively, bridging α8 and β9. Notably, Arg100_CDRH3_ was observed in training antibodies but was not found in the most similar training antibody (VH distance = 12) despite having a highly similar CDRH3 (94.4% identity). CDRH1 and CDRH2 make polar and hydrophobic contacts within and proximal to the α8 helix, including two salt bridges formed between Arg32_CDRH1_ and Glu378_F_ and Asp58_CDRH2_ and Lys390_F_. The RSV-3301 light chain also buries surface area on F between α8 and β2, primarily through hydrophobic contacts mediated by Tyr32_CDRL1_ and Tyr92_CDRL3_. Additionally, CDRL1 and LFR3 contact residues within β22, extending the RSV-3301 epitope into antigenic Site IV.

Together, the structural characterization of these two antibodies demonstrates that MAGE generates antibodies with diverse binding properties. Not only do RSV-2245 and RSV-3301 target different binding sites of the RSV-A F protein, but these two antibodies display different binding properties. RSV-2245 contains binding residues distributed across both the heavy and light chains, whereas RSV-3301 binding is dominated by interactions within CDRH3. Although both antibodies contained MAGE-generated mutations in key binding residues, there were many mutations introduced into framework regions that did not interface with the antigen surface. To test the impact of these non-germline mutations, we reverted the VH genes to germline and tested for binding by ELISA, with the germline-reverted RSV-3301 showing substantial reduction in binding by ELISA, while germline-reverted RSV-2245 retained comparable binding to its fully mutated form (**Fig. S8a-b**). Further, BLI was used to characterize the binding of RSV-2245 Fab and RSV-3301 Fab to immobilized RSV-A F (**Fig. S8c-d**). For RSV-2245, a 1:1 binding model was used to determine binding affinity *(K*_D_ = 1.5×10^-7^ M). Due to suspected heterogeneity in the epitope targeted by RSV-3301, these data were fit to a heterogeneous ligand model to determine two *K*_D_ values (*K*_D*1*_ = 6.7×10^-6^ M and *K*_D*2*_ *=* 4.5×10^-9^ M). Together, these results show that MAGE can generate antibodies with a variety of SHM changes in different regions of the antibody sequence and with differing impacts on antigen recognition and binding affinity.

## Discussion

In this work, we aimed to develop a purely sequence-based model capable of generating paired heavy-light chain antibody sequences with prompt-specific binding. Once trained, the MAGE model presented here requires no template antibody or protein structural information. When prompted with an antigen amino acid sequence, MAGE produces full human variable heavy and light chains, including novel designs with changes from germline sequence introduced throughout the entire variable sequences. Our results confirm that generative LLM models like MAGE are capable of the complex task of generating full paired heavy and light chain antibody sequences, demonstrating validated binding against RBD, H5 hemagglutinin, and RSV-A prefusion F. MAGE-generated antibodies showed diverse sequence characteristics and binding properties, including potent neutralization for a subset of the binding antibodies designed against each antigen. While MAGE is not conditioned on neutralization, this demonstrates the functionality of these antibodies and validates the ability of MAGE to produce useful, clinically relevant antibodies in the context of therapeutic discovery. For RBD and RSV-A, a subset of validated, target-specific designs were selected with no bias towards known antibodies, demonstrating design of potently neutralizing antibodies without the need for a starting template antibody sequence or structure. The design of neutralizing antibodies against H5/TX/24 hemagglutinin demonstrates zero-shot learning capabilities, where MAGE was able to generate antibodies against an unseen influenza strain by training on previously characterized antibodies with specificity against a related, but divergent H5N1 strain. This demonstrates a realistic use-case for this approach, where MAGE could be used to generate antibodies against an emerging health threat more rapidly than traditional antibody discovery methods that would rely on access to specialized biological materials (e.g. blood samples or antigen protein).

The antibodies designed and characterized here display a range of sequence characteristics, including differential gene usage, CDR properties, and levels of SHM. While a subset of the validated binding antibodies have CDRH3s that are similar to those seen in training, it is well-established that individual amino acid substitutions can disrupt antibody-antigen binding(*41, 42*), even within non-interfacing framework regions(*43, 44*). As such, the ability of the model presented here to generate binding – and in some cases potently neutralizing – antibody sequences highlights the utility of generative algorithms in creating solutions that differ from those seen in the training data while retaining antibody-antigen recognition properties. In addition to designs with low numbers of edits introduced to training antibodies, we also validated binding for more novel antibodies with >20 total amino acid differences to the most similar training examples (RSV-2245 and RSV-3301). Structural characterization of these antibodies showed that they target different sites on RSV F with different modes of binding which utilize residues not found in the closest training antibody matches. Additionally, the Site I epitope targeted by RSV-3301 is not well characterized and, to our knowledge, this is the first structure showing a human antibody targeting this epitope in prefusion F(*45*).

We emphasize that MAGE is not restricted to redesigning existing antibodies, rather it is able to sample the distribution of known binding sequences to learn the complex sequence features associated with antigen-binding specificity and then generate a pool of diverse antibodies that is highly enriched for binding antibodies, providing candidates for further characterization, down-selection, and development. In this study we have only sampled a fraction of this sequence space for validation but envision that this candidate pool could be further mined to find antibodies with desired properties that have not yet been explored.

In this work, MAGE was validated against viral antigen targets as a proof-of-concept. However, data generation methods are constantly improving, and large-scale efforts using high-throughput methods such as LIBRA-seq could soon yield datasets of sufficient scale for training such models to efficiently generate antibodies against diverse antigen targets beyond what is included in the training datasets. The development of these datasets, along with the subsequent experimental validation of generated antibodies which can be incorporated into training data, will enable iterative improvement of MAGE. Since applications of LLMs in other fields have shown evidence of generalization(*46–48*), we anticipate that, provided enough data, models such as MAGE could be capable of learning the more general rules of residue-level interactions that govern antibody-antigen binding with the capability to generate antibodies against completely unseen targets. Such approaches will have the potential to revolutionize the field of antibody discovery, but the generalizability of such models is yet to be proven in this context. Despite these limitations, the model described here presents the first example of an LLM capable of antigen-specific paired heavy-light chain antibody sequence generation and provides a promising glimpse into the future of AI-accelerated antibody discovery.

## Acknowledgements

We thank all members of the Georgiev laboratory for their support and feedback. We thank Ben Murrell for feedback on the manuscript. We thank Vito Quaranta and Darren Tyson for providing computational resources through the Quantitative Systems Biology Center. We thank the Vanderbilt Technologies for Advanced Genomics Core (VANTAGE), which is supported in part by CTSA (5UL1 RR024975-03), for providing technical assistance with library production and sequencing. We would like to thank Mark Connors for providing HIV-1 PBMC samples. Biorender was used for creating schematic figures.

## Funding

This research was funded, in part, by the Advanced Research Projects Agency for Health (ARPA-H). The views and conclusions contained in this document are those of the authors and should not be interpreted as representing the official policies, either expressed or implied, of the U.S. Government.

G. Harold and Leila Y. Mathers Charitable Foundation grant MF-2107-01851 (ISG)

National Institutes of Health grant R01AI175245 (ISG)

National Institutes of Health grant R01 AI152693 (ISG)

Swedish Research Council 2018-02381 (Ben Murrell)

Swedish Research Council 2023-02516 (Ben Murrell)

Welch Foundation grant F-0003-19620604 (JSM)

## Author contributions

Conceptualization: PTW, TMM, ISG

Methodology: PTW ISG

Investigation: PTW, NVJ, AKJ, JRK, TMM, GH, JGAS, TMR, GJ, MJV, SH, LV, DJS, RAG, SFA, MK, GAS, FP, CMH

Project administration: GH, JL, HYC, GAS, TMR

Supervision: JSM, AAA-S, ISG

Writing – original draft: PTW, ISG

Writing – review & editing: PTW, NVJ, AKJ, JRK, TMM, GH, JGAS, TMR, GJ, MJV, SH, LV, DJS, RAG, SFA, MK, GAS, FP, CMH, JSM, AAA-S, ISG

## Competing interests

I.S.G. is a cofounder of AbSeek Bio. P.T.W and I.S.G. are listed as inventors on patents filed describing the pipeline presented here for the fine-tuning of LLMs for antigen-specific antibody generation. The Georgiev laboratory has received unrelated funding from Takeda and Merck. Dr. Chu has consulted for Bill and Melinda Gates Foundation and Ellume, and has served on advisory boards for Vir, Merck and Abbvie; she has received research funding from Gates Ventures, and support and reagents from Ellume and Cepheid outside of the submitted work.

## Data and materials availability

Data will be made available upon publication. Raw sequencing reads from LIBRA-seq will be uploaded to the National Library of Medicine Sequence Read Archive (SRA). The Cryo-EM structure of Fabs RSV-2245 and RSV-3301 in complex with RSV-A prefusion F will uploaded to the PDB. The full curated database will be provided along with the fine-tuned model on HuggingFace.

## Materials and Methods

### Antibody generation and basic filtering

For generation of antibodies against an antigen target, the fine-tuned model was prompted with the entire antigen amino acid sequence. The antibody sequences were annotated using ANARCI(*55*) with IMGT(*26*) numbering. If a sequence had either a heavy or light chain that was not recognized as a human variable chain by ANARCI, it was discarded, along with any sequences missing framework or CDR regions following alignment. Heavy chains or light chains shorter than 100 amino acids were also discarded, although few sequences under these lengths survived the previous filtering steps. Heavy chains which were identical to a training example were discarded, although there was only one occurrence of this in the sequences generated against SARS-CoV-2 RBD. Finally, antibodies were assessed for mutational load based on identity to germline variable genes, and humanness using the BioPhi OASis software.

Perplexity was calculated by performing a backward pass through the trained model, with the model in evaluation mode, to calculate the exponential of the average negative log-likelihood for each generated sequence.

### Antibody selection for experimental validation of RBD antibodies

In addition to the basic filtering outlined above, additional criteria were applied to select candidates for experimental validation. For the heavy variable gene, only sequences with >85% identity were retained. For humanness, sequences below 70^th^ percentile were discarded based on the recommended threshold for OASis(*27*). In order to test novel sequences generated by the model, antibodies with a CDRH3 identical to any training example (n = 72), or a VH germline identity of 100% (n = 175) were removed, leaving a selection pool of 732 antibodies. In addition, any sequences with both >90% VH identity and >90% CDRH3 identity to CoV-AbDab RBD binding antibody sequences were removed. Following these filtering steps, the following automated selected pipeline was applied:

1. From the top 5 most frequently generated VH genes, select sequences with 20^th^ and 80^th^ percentile VH identities.
2. Select top 5 sequences by rank of maximum CDRH3 identity to known binding antibodies.
3. Select top 5 sequences by rank of maximum VH identity to know binding antibodies.

Together, these three selection steps yield 20 antibodies per antigen target. Antibodies from Step 1 represent selection independent of known binding antibodies to avoid any bias, while antibodies from Steps 2 and 3 yield testing antibodies with high similarity to known binding antibodies.

### Antibody selection for experimental validation of RSV antibodies

MAGE was prompted using the RSV-A Fusion glycoprotein F0 (UniProt(*52*) entry P03420) amino acid sequence to generate 10,000 sequences for down selection and validation. Basic filtering was applied as described above, along with filtering based on perplexity (PPL < 1.5), heavy chain germline identity (percent VH identity < 98%), and sequence identity to training antibodies (maximum CDRH3 identity to any training antibody ≤ 95%). Due to the overrepresentation of CoV-specific antibodies seen in training, the remaining generated sequences were compared to CoV-AbDab antibodies to remove sequences with CDRH3s similar to CoV-specific antibodies (CDRH3 percent identity > 70%).

Following filtering, antibodies were selected for validation based on three separate criteria groups. First, an unbiased group was selected by clustering generated CDRH3s using hierarchical clustering based on a Levenshtein identity matrix with a maximum identity distance of 20% within each cluster. One sequence was then randomly sampled from each of the top 10 largest clusters, for a total of 10 unbiased sequences selected for validation. For the unbiased group, we selected generated sequences with CDRH3 identity ≥75% and equal CDRH3 length compared RSV-A-specific training antibodies. From these matches, 10 antibodies were randomly selected from unique CDRH3 clusters. Finally, three generated antibodies with CDRH3 identity ≥70% to RSV-A-specific training antibodies and CDRH3 identity ≥60% to MPV-A-specific training antibodies were selected. In total, 23 antibodies were selected for experimental validation.

### Antibody selection for experimental validation of H5N1 antibodies

MAGE was prompted using the highly pathogenic avian influenza virus H5/TX/24 hemagglutinin sequence (Strain A/Texas/37/2024, GenBank accession number PP577943.1(*34*)) amino acid sequence as a prompt to generate 1,000 sequences for down selection and validation. Basic filtering was applied as described above, along with filtering based on heavy chain germline identity (percent VH identity < 100%). Antibodies were then selected for validation based on CDRH3 Levenshtein identity to H5-specific training antibodies. Generated sequences were randomly selected from four CDRH3 identity bins: [80% - 85%) (n = 3), [85% - 90%) (n =6), [90% - 95%) (n = 6), and [95% - 99%) (n = 3) for a total of 18 antibodies. Since this exact flu strain sequence was not seen in training, no unbiased group was selected for testing.

### Antibody expression and purification

Variable genes were synthesized as cDNA and were inserted into bi-cistronic plasmids encoding for the constant regions of the heavy chain and either the kappa or lambda light chain, for each antibody (Twist BioScience). DH5α cells were transformed with the antibody DNA, and the resulting ampicillin resistant colonies were grown in LB broth. Plasmid DNA was isolated from the bacterial cultures using a plasmid purification kit (Qiagen). The purified antibody DNA was transfected into Expi293F cells using ExpiFectamine transfection reagent (Thermo-Fisher Scientific), and antibodies transiently expressed in FreeStyle F17 expression media (Thermo-Fisher) supplemented 0.1% Pluronic Acid F-68 and 20% 4 mM L-glutamine. Cells were cultured at 8% CO_2_ saturation and 37°C with shaking. Cells were collected five days post transfection and centrifuged at a minimum of 5,000 rpm for 20 minutes. Filtered supernatant (Nalgene Rapid Flow Disposable Filter Units with PES membrane 0.45 or 0.22 μm) was purified over protein A equilibrated with PBS. Antibodies were eluted from the column with 100 mM glycine HCl at pH 2.7 directly into a 1:10 volume of 1 M Tris-HCl pH 8 and then buffer exchanged into PBS for storage at 4°C.

### Enzyme-linked immunosorbent assay (ELISA)

Recombinant antigen (SARS-CoV-2 Index RBD, SARS-CoV-2 Index S, XBB.1 spike, SARS-CoV-1 S) was plated at 2 ug/mL overnight at 4°C. The next day, plates were washed three times with PBS supplemented with 0.05% Tween20 (PBS-T) and coated with 5% bovine serum albumin (BSA) in PBS-T. Following a one-hour incubation at room temperature, the plates were washed three times with PBS-T. Primary antibodies diluted in 1% BSA in PBS-T were then added to the plates, starting at 10 μg/mL with a serial 1:5 dilution, followed by a one-hour incubation at room temperature. Plates were then washed three times in PBS-T before adding secondary antibody, goat anti-human IgG conjugated to peroxidase, at 1:10,000 dilution in 1% BSA in PBS-T followed by a one-hour incubation at room temperature. Plates were washed for a final three times with PBS-T and then developed by adding TMB substrate to each well. Plates were incubated at room temperature for five minutes, and then 1 N sulfuric acid was added to stop the reaction. Plates were read at 450 nm. ELISAs were performed in technical and biological duplicate. The area under the curve (AUC) values were calculated using GraphPad Prism 9.5.0 to fit a 4-parameter log(agonist) vs. response curve.

### Antigen expression and purification

For the different binding experiments, SARS-CoV-2 Index S RBD (2019-nCoV) was purchased from Sino Biological catalog number (40592-VNAH) while SARS-CoV-2 S Hexapro Index strain, SARS-CoV-2 S XBB.1, and SARS-CoV-1 S were expressed in Expi293F cells by transient transfection in FreeStyle F17 expression media (Thermo-Fisher) supplemented to a final concentration of 0.1% Pluronic Acid F-68 and 20% 4 mM L-glutamine using ExpiFectamine transfection reagent (Thermo-Fisher) cultured for 4-7 days at 8% CO_2_ saturation and 37°C with shaking. After transfection, cultures were centrifuged at 5000 rpm for 20 minutes. Filtered supernatant (Nalgene Rapid Flow Disposable Filter Units with PES membrane 0.45 or 0.22 μm), was run slowly over equilibrated, 1 mL pre-packed StrepTrap XT column (Cytiva Life Sciences). The column was washed with 15 mL of binding buffer (100 mM Tris-HCl, 150 mM NaCl, 1 mM EDTA, pH 8.0), and purified protein was eluted from the column with 10 mL of binding buffer supplemented with 2.5 mM desthiobiotin. Protein was concentrated, buffer exchanged into PBS and run on a Superose 6 Increase 10/300 GL on the AKTA FPLC system. Peaks corresponding to trimeric species were identified based on elution volume and SDS-PAGE of elution fractions. Fractions containing pure spike were pooled.

#### Biolayer interferometry

BLI experiments were performed using an OctetRED96e instrument (Sartorius) at 21°C and a shaking speed of 1000 rpm. For the RBD-binding antibodies, purified SARS-CoV-2 Wuhan-Hu-1 RBD-SD1 (residues 319–591) containing a C-terminal 8xHis tag was immobilized to Ni-NTA sensortips (Sartorius) to a response level of approximately 1.5 nm in HBS-P buffer (10 mM HEPES pH 7.4, 150 mM NaCl, 0.005% v/v Surfactant P20) with 20 mM imidazole and 0.1% w/v BSA added. After a 60 s baseline step, immobilized RBD-SD1 was dipped into wells containing 2-fold serial dilutions of IgG ranging in concentration from 32 to 0.5 nM (RBD-159, RBD-238, RBD-409, RBD-839, and RBD-951) or 1024–16 nM (RBD-446) to measure association. 1.5-fold dilutions ranging from 1024 to 90 nM of RBD-61 and a combination of 1.5-fold (1024–303 nM) and 2-fold (303–38 nM) dilutions of RBD-413 were used to optimize the dynamic range of the binding curves for those antibodies. Dissociation was measured by dipping sensortips into wells containing buffer only. Data were reference subtracted and kinetics were calculated (high-affinity antibodies only) by fitting curves to a 1:2 bivalent analyte model using the Octet Data Analysis Software v11.1.

Binding specificity was measured by immobilizing 8xHis-tagged SARS-CoV-2 Wuhan-Hu-1 RBD-SD1 or 8xHis-tagged prefusion-stabilized RSV F trimer (DS-Cav1(*30*)) to Ni-NTA sensortips to a response level of approximately 1.5 nm in the buffer described above. Immobilized antigen was then dipped into wells containing the anti-RBD IgG of interest (4, 16, or 512 nM antibody for immobilized RBD-SD1 and 512 nM antibody for immobilized RSV F). Immobilized RSV F was also dipped into wells containing only buffer to observe baseline signal drift.

For the RSV F-binding antibodies, purified 8xHis-tagged prefusion-stabilized RSV F trimer (DS-Cav1) was immobilized to Ni-NTA sensortips to a response level of approximately 0.8 nm in HBS-P buffer with 20 mM imidazole and 0.1% w/v BSA added. After a 60 s baseline step, immobilized Ds-Cav1 was dipped into wells containing 2-fold serial dilutions of Fab ranging in concentration from 640 to 10 nM (RSV-2245) or 5 to 0.78 μM (RSV-3301) to measure association. Dissociation was measured by dipping sensortips into wells containing buffer only. Data were reference subtracted and kinetics were calculated by fitting curves to a 1:1 (RSV-2245) or heterogeneous ligand (RSV-3301) model using the Octet Data Analysis Software v11.1.

### SARS-CoV-2 Pseudovirus Neutralization Assay

Pseudovirus neutralization assays were performed as previously described(*56*). Briefly, spike-pseudotyped lentiviruses were produced by the co-transfection of HEK293T cells with respective spike variant plasmids, together with an HIV gag-pol packaging plasmid (Addgene #8455) and a firefly luciferase encoding transfer plasmid (Addgene #170674). Transfections were performed using polyethylenimine. Pseudoviruses titrated to produce approximately 100,000 RLU were incubated with 8 serial 3-fold dilutions for 1 hour at 37°C in black-walled 96-well plates. 10,000 HEK293T-ACE2 cells were then added to each well, and plates were incubated at 37°C. Luminescence was measured approximately 48 hours later on a GloMax Navigator Luminometer (Promega) using Bright-Glo luciferase substrate (Promega) as per the manufacturer’s recommendations. Neutralization was calculated relative to the average of 8 control wells infected in the absence of antibody. IC_50_ values were calculated by fitting a four-parameter logistic curve and interpolating the concentration at which there is 50% neutralization, using Prism v10.1.0 (GraphPad Software).

### RSV Neutralization Assay

RSV neutralization assays were performed similarly to previously described protocols (PMID: 37403896). In brief, Vero cells were seeded the day before the assay at a density of 2×10^4^ cells per well in a 96-well plate in high glucose DMEM supplemented with L-glutamine and 10% FBS. Monoclonal antibodies were 2-fold serially diluted starting from a concentration of 50 μg/mL in DMEM supplemented with 2% FBS and each antibody dilution was mixed with an equal volume of the same medium containing 100 TCID50 of RSV virus strain A2 (cat n. NR-52018, BEI Resources) and incubated for 1 hour at 37°C, 5% CO_2_. Uninfected and infected cell wells without the antibody were also included as controls. After the incubation, the antibody-virus mixtures were added to the cells and plates were incubated at 37°C, 5% CO_2_ for 72 hours. Plates were then washed with PBS and cells fixed with cold 80% v/v acetone in PBS for 10 minutes at RT. After the incubation, the plates were emptied and washed 3 times with wash buffer (PBS + 0.3% Tween20). Primary mouse anti-RSV F antibody (cat n. MCA490, Bio-Rad) diluted at 1:1,000 in blocking buffer (wash buffer + 7.5% BSA) was then added to the plates and incubated for 1 hour at RT. Following the incubation, the plates were washed 3 times with wash buffer and secondary goat anti-mouse IgG human adsorbed HRP-conjugated secondary antibody (cat. n. 1030-05, Southern Biotech) diluted 1:1,000 was added and incubated for 1 hour in the dark at RT. Plates were then washed 5 times with wash buffer and freshly prepared o-Phenylenediamine dihydrochloride (OPD) substrate added and incubated for 3-5 minutes at RT. Reaction was stopped by adding 2N H_2_SO_4_ and absorbance read at 490 nm using a PowerWaveXS plate reader (BioTek).

### Influenza reporter virus neutralization assay

Generation of the replication-restricted reporter (R3ΔPB1) H1N1 virus (A/Michigan/45/2015) as well as rewired R3ΔPB1 (R4ΔPB1) H5N1 virus (A/Vietnam/1203/2004) is described elsewhere(*57*). R4ΔPB1 H5N1 A/Texas/37/2024 virus was prepared similarly. Briefly, to generate the R3/R4ΔPB1 viruses the viral genomic RNA encoding functional PB1 was replaced with a gene encoding the fluorescent protein (TdKatushka2), and the R3/R4ΔPB1 viruses were rescued by reverse genetics and propagated in the complementary cell line which expresses PB1 constitutively. Each R3/R4ΔPB1 virus stock was titrated by determining the fluorescent units per mL (FU ml^-1^) prior to use in the experiments. For virus titration, serial dilutions of virus stock in OptiMEM were mixed with pre-washed MDCK-SIAT1-PB1 cells (8 × 10^5^ cells/ml) and incubated in a 384-well plate in quadruplicate (25 µl well^-1^). Plates were incubated for 18–26 h at 37°C with 5% CO_2_ humidified atmosphere. After incubation, fluorescent cells were counted by using a Celigo Image Cytometer (Nexcelom) with a customized red filter for detecting TdKatushka2. For the microneutralization assay, serially diluted antibodies were prepared in OptiMEM and mixed with an equal volume of R3/R4ΔPB1 virus (∼8 × 10^4^ FU ml^-1^) in OptiMEM. After incubation at 37°C and 5% CO_2_ humidified atmosphere for 1 h, pre-washed MDCK-SIAT1-PB1 cells (8 × 10^5^ cells well^-1^) were added to the antibody-virus mixtures and transferred to 384-well plates in quadruplicate (25 µl well^-1^). Plates were incubated and counted as described above. Target virus control range for this assay is 500 to 2,000 FU per well, and cell-only control is acceptable up to 30 FU per well. The percent neutralization was calculated for each well by constraining the virus control (virus plus cells) as 0% neutralization and the cell-only control (no virus) as 100% neutralization. A 7-point neutralization curve was plotted against antibody concentration for each sample, and a four-parameter nonlinear fit was generated using Prism (GraphPad) to calculate the 50% (IC_50_) inhibitory concentrations.

### LIBRA-seq Experiments

For the different LIBRA-seq experiments, a total of 24 proteins were expressed as recombinant soluble antigens. Influenza, parainfluenza, coronavirus, RSV post fusion, hMPV post fusion, and HIV-1 antigens were expressed as described above and then purified over the appropriate affinity column at 4°C.

Recombinant hemagglutinin (HA) proteins all contained the HA ectodomain with a point mutation at the sialic acid-binding site (Y98F), a T4 fibritin foldon trimerization domain, and a hexahistidine-tag. HAs were purified by metal affinity chromatography. Parainfluenza virus type 3 prefusion stabilized F ectodomain (PDB: 6MJZ) was purified by nickel affinity chromatography. SARS-CoV-2 S XBB.1, BQ.1.1, SARS-CoV-1 S, HCoV-OC43 S, HCoV-HKU1-S-2P, RSV post fusion, and hMPV post fusion were purified over pre-packed StrepTrap XT column (Cytiva Life Sciences), as described above. Single chain HIV-1 gp140 SOSIP variant strain BG505(*58*) was purified over agarose bound Galanthus nivalis lectin (Vector Laboratories cat no. AL-1243-5). Methods have been previously described.(*59*)

Previously described hMPV F A1(NL/1/00) and B2(TN99-419) antigens were expressed in FreeStyle 293-F cells by transient transfection in FreeStyle 293 expression media (Thermo-Fisher). Cells were co-transfected at a 4:1 ratio of plasmids encoding human metapneumovirus F and furin, respectively, using polyethylenimine (PEI). Three hours post-transfection, media was supplemented to a final concentration of 0.1% (v/v) Pluronic Acid F-68. After culturing for 6 days at 37°C and 8% CO_2_ saturation, filtered supernatant was concentrated and buffer exchanged to PBS using tangential flow filtration. Samples were then run over a gravity-flow affinity column at room temperature. Previously described RSV F (DS-Cav1) A2 and B9320 antigens were expressed similarly but did not include the Pluronic F-68 supplementation step. Stabilized ectodomains of hMPV F subtypes A1 and B2(*60*), as well as RSV strains A2(*30, 61*) and B9320 F (DS-Cav1(*62*)), were purified over Strep-Tactin Sepharose resin (IBA Lifesciences) in a gravity column.

CHDC, SYD_2012, and GII.17 P domains were recombinantly expressed and purified as previously described(*63*).

All proteins were quantified using UV/vis spectroscopy. Antigenicity of proteins was characterized by ELISA with known monoclonal antibodies specific for that antigen. Proteins were frozen and stored at −80°C until use.

### Donor peripheral blood mononuclear cell (PBMCs) samples

Healthy peripheral blood mononuclear cell (PBMC) samples were purchased from StemCell Technologies. SARS-CoV-2 PBMCs were collected from individuals with SARS-CoV-2 infection, 60 days post symptom onset during May-June 2020. Influenza vaccination PBMCs were collected from individuals 28 days following vaccination with the 2021-2022 quadrivalent flu vaccine. HIV-1 PBMCS were collected between 2007-2013 from individuals with confirmed HIV-1 status.

### Conjugation of oligonucleotide barcodes to antigens for LIBRA-seq

For each antigen, a unique DNA barcode was directly conjugated to the antigen using a SoluLINK Protein-Oligonucleotide Conjugation kit (TriLink, S-9011) according to manufacturer’s protocol.

### Biotinylation of antigens for LIBRA-seq

Protein antigens were biotinylated using EZ-link Sulfo-NHS-Biotin No-Weigh kit (Thermo Fisher) according to manufacturer’s instructions. A 50:1 biotin-to-protein molar ratio was used for all reactions.

### Flow cytometry enrichment of antigen-specific B cells

For a given sample, cell mixtures were stained and mixed with fluorescently labeled DNA-barcoded antigens and other antibodies, and then sorted using fluorescence activated cell sorting (FACS). Cells were counted, washed with DPBS supplemented with 0.1% Bovine serum albumin (BSA), and resuspended in DPBS-BSA to be stained with the following cell markers: Ghost Red 780, CD14-APCCy7, CD3-FITC, CD19-BV711, and IgG-PECy5. Additionally, antigen-oligo conjugates were added to the stain. Following a 30-minute incubation in the dark on ice, the cells were washed three times with DPBS-BSA then incubated for 15 minutes in the dark on ice with Streptavidin-PE label cells with bound antigen. Cells were then resuspended in DPBS-BSA and sorted on the cell sorter. Antigen positive cells were bulk sorted and then delivered to the Vanderbilt VANTAGE sequencing core at an appropriate target concentration for 10X Genomics library preparation and subsequent sequencing. FACS data were analyzed using FlowJo.

### Sequence processing and bioinformatics analysis for LIBRA-seq

We followed our established pipeline(*21*), which takes paired-end FASTQ files of oligonucleotide libraries as input, to process and annotate reads for cell barcodes, unique molecular identifiers (UMIs) and antigen barcodes, resulting in a cell barcode-antigen barcode UMI count matrix. B cell receptor contigs were processed using CellRanger 3.1.0 (10x Genomics) and GRCh38 Human V(D)J 7.0.0 as reference, while the antigen barcode libraries were also processed using CellRanger (10x Genomics). The cell barcodes that overlapped between the two libraries formed the basis of the subsequent analysis. Cell barcodes that had only non-functional heavy chain sequences as well as cells with multiple functional heavy chain sequences and/or multiple functional light chain sequences, were eliminated, reasoning that these may be multiplets. We also aligned the B cell receptor contigs to IMGT reference genes using HighV-Quest(*64*). The annotated sequences were then combined with an antigen barcode UMI count matrix. Finally, we determined the LIBRA-seq score (LSS) for each antigen in the library for every cell as previously described(*21*). Binding was defined using a conservative threshold of LSS ≥ 2, based on validation results from previous LIBRA-seq studies. Cells which bound to multiple antigens from different viral families were filtered out to remove polyreactive BCRs, along with any cells from non-HIV donors which bound HIV antigens.

### Cryo-EM sample prep and data collection

RSV F (prefusion-stabilized, PR-DM(*38*) was mixed to a final concentration of 2.5 mg/mL with 1.5X molar excess of Fabs RSV-2245 and RSV-3301 in buffer containing 2 mM Tris pH 7.5, 200 mM NaCl, 0.02% NaN_3_. The complex was incubated for 30 minutes at 4 °C before adding 10X CMC CHAPS (VitroEase™ Buffer Screening Kit, Thermo Fisher) to a final concentration of 0.1X CMC. Immediately following the addition of CHAPS, 3.5 μL of sample was applied to C-flat 1.2/1.3 300 mesh grids (Electron Microscopy Sciences) that had been glow discharged using a PELCO easiGlow (Ted Pella) for 30 seconds at a current of 20 mA. Using a Vitrobot Mark IV (Thermo Fisher), a blot force of 1 was applied for 9 s to blot away excess liquid before plunge-freezing into liquid ethane. Samples were blotted in 100% humidity at 4 °C.

1,561 movies were collected from a single grid using a Glacios TEM (Thermo Fisher) equipped with a Falcon 4 detector (Thermo Fisher), with the stage tilted to 30°. All movies were collected using SerialEM v4.0.10 automation software(*65*). Particles were imaged at a calibrated magnification of 0.933 Å/pixel, with an exposure of 2.5 eps for 17s for a total exposure of 49 e/Å2. Additional details about data collection parameters can be found in Supplementary Table 3.

### Cryo-EM processing and structure building

Motion correction, CTF estimation, particle picking, and preliminary 2D classification were performed using cryoSPARC v4.6.0 live processing(*66*) (Supplementary Figure 1 workflow). An initial ab initio reconstruction of four classes was performed during live processing using 123,151 particles. Once data collection was completed, a final iteration of 2D class averaging distributed 610,184 particles into 80 classes using an uncertainty factor of 1 and a batchsize of 300 for 25 iterations. From that, 338,721 particles were selected and carried into a heterogeneous refinement of the four volumes that resulted from the initial ab initio reconstruction. Particles from the highest quality class were used for homogenous refinement of the best volume with applied C3 symmetry. To address remaining particle heterogeneity, 210,844 particles (after re-extraction and duplicate removal) were sorted into four classes by performing another Ab initio reconstruction, followed by heterogeneous refinement of the four classes using all particles. From this, 140,634 particles were taken from the best class and used for a final non-uniform refinement with applied C3 symmetry and with refined per-particle defocus and per-group CTF parameters(*67*). To improve map quality, the refinement volumes were processed using DeepEMhancer(*68*) within cryoSPARC(*66*) [cite]. An initial model of the complex was generated using AlphaFold 3 (https://alphafoldserver.com) by inputting sequences (separately) for RSV F1 and F2, 2245 VH and VL, and 3301 VH and VL(*69*). The highest confidence output model was docked into the refined volume via ChimeraX v1.8(*70*). The structure was iteratively refined and completed using a combination of Phenix v1.21.2(*71*), Coot v0.9.2(*72*), and ISOLDE v1.8(*73*).

## Supplementary Data and Figures

**Fig. S2.**
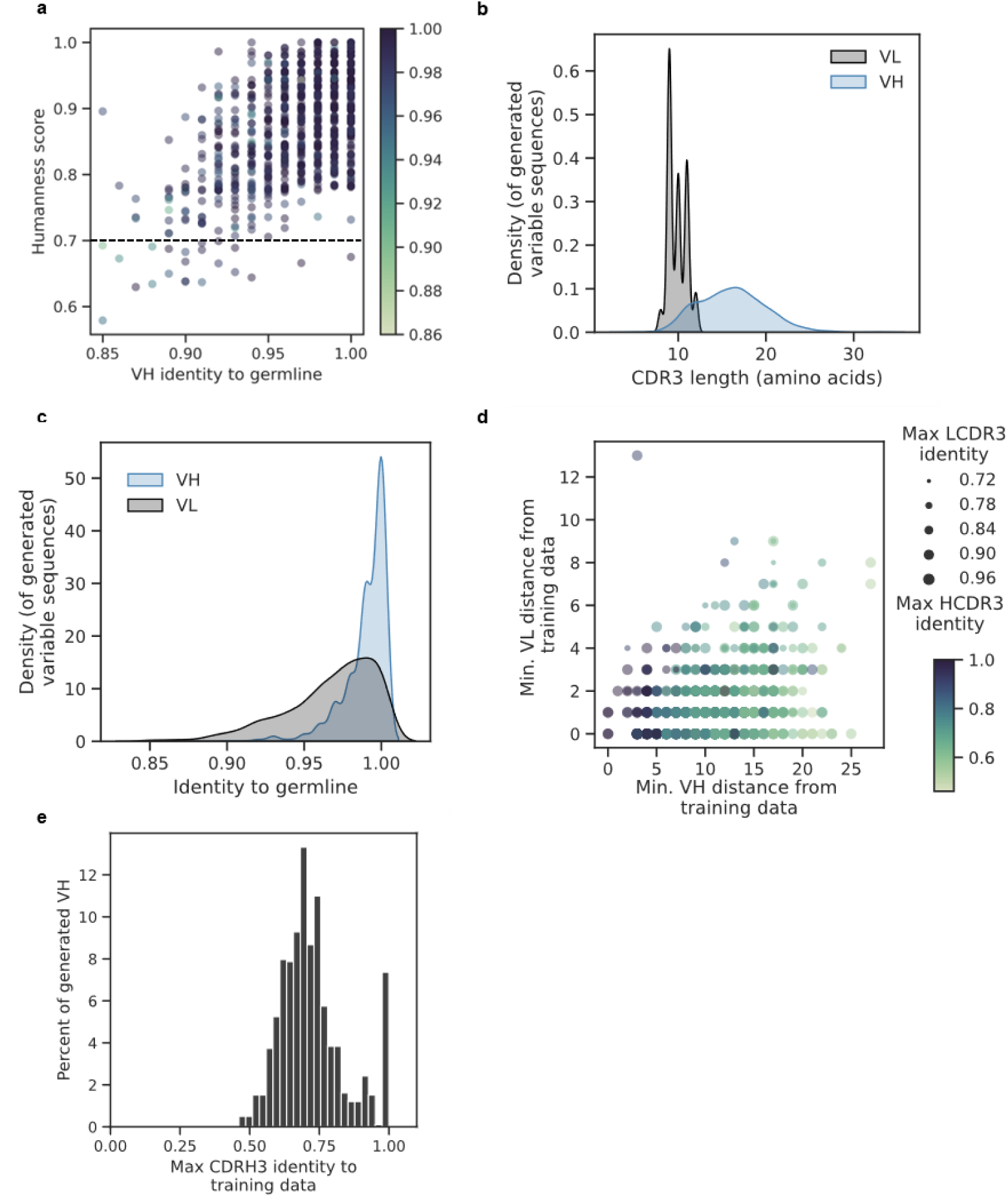
A) Scatterplot showing relationship between VH identity and the OASis humanness score for antibodies generated against RBD. Dotted line represents the threshold used. B) Distributions of CDR3 length for heavy and light variable sequences in generated antibodies. C) Distributions of percent identity to germline for variable heavy (VH) and light (VL) chains in antibodies generated against RBD. D) Based on the comparison in C, the distance to the closest training example for the generated VH and VL sequences is shown. Size of the points represents the maximum LCDR3 to any training sequence, and color represents the maximum HCDR3 to any training sequence. E) Distribution of the maximum CDRH3 identity between each generated antibody and the training data.

**Fig. S3.**
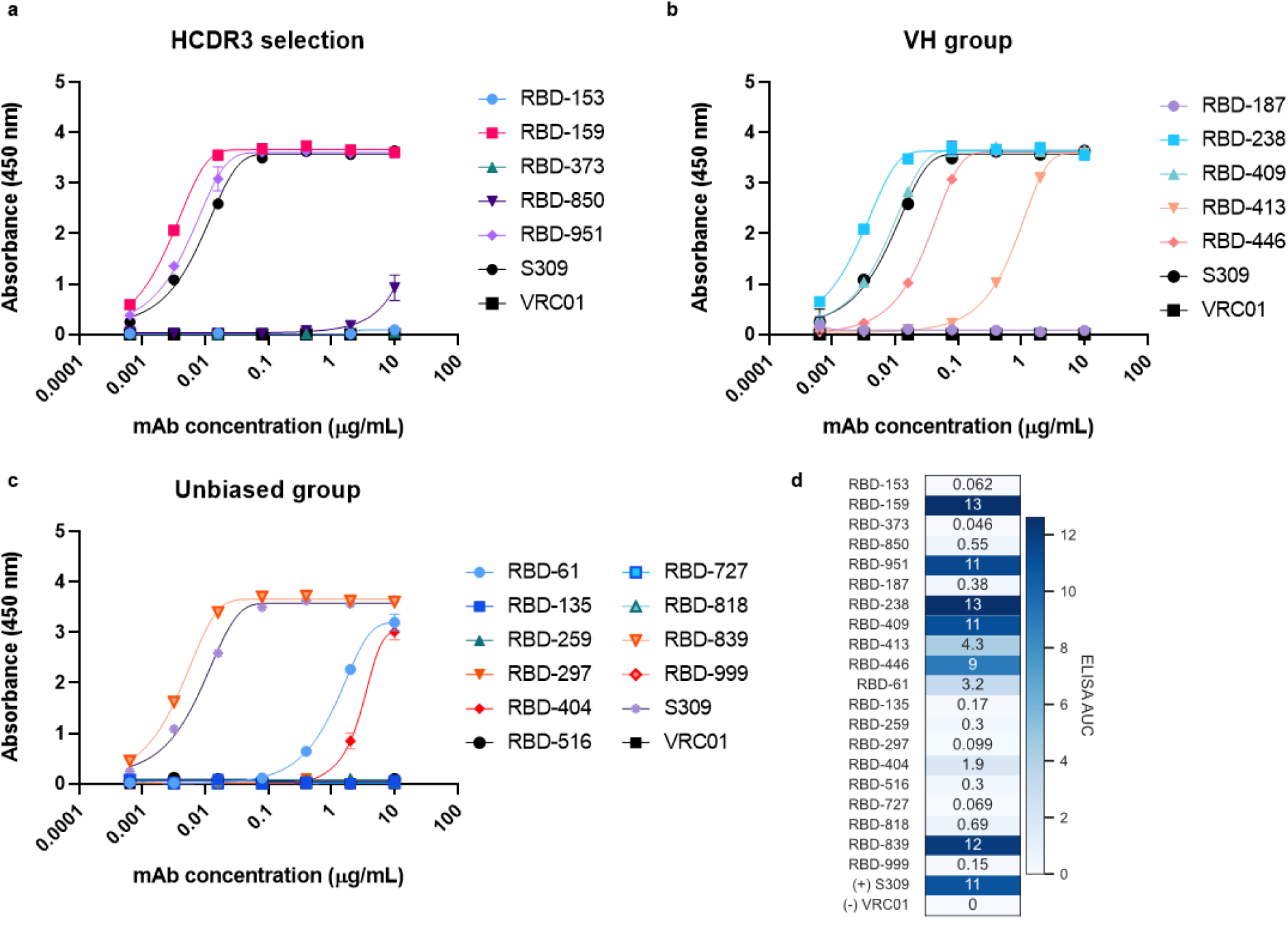
ELISA dilution curves for A) HCDR3 identity selection group, B) VH identity selection group, and C) unbiased selection group. Positive control SARS-CoV-2 RBD binding antibody (S309) and negative control HIV-1 specific antibody (VRC01) are shown in black. D) ELISA AUCs from curve fit to dilution series. Same as Fig. 2c, but with numbers shown on heatmap.

**Fig. S4.**
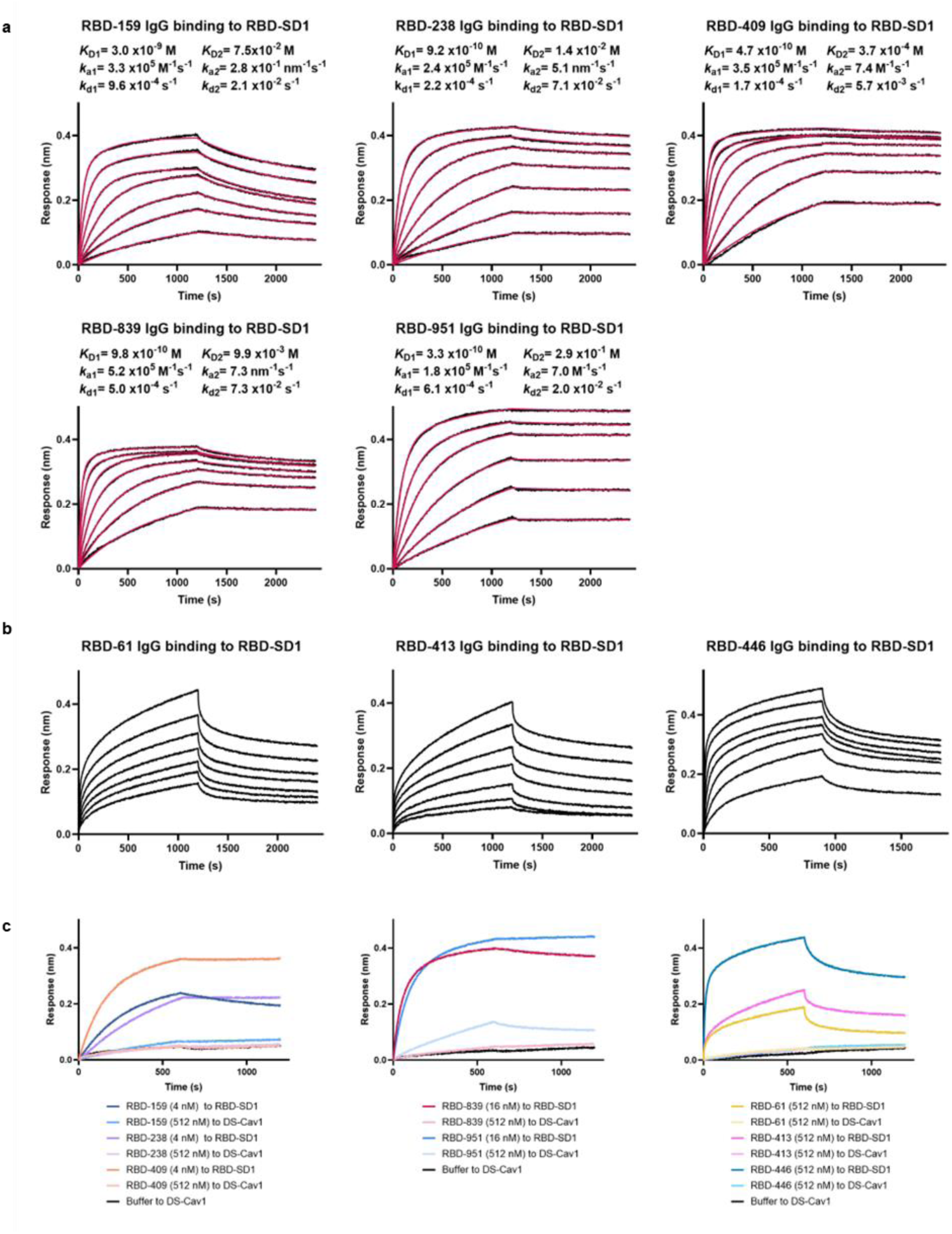
A) BLI sensorgrams (also shown in Figure 2E) for the association and dissociation kinetics of high-affinity antibodies binding to immobilized SARS-CoV-2 RBD-SD1. Data (black) were fit to a 1:2 bivalent analyte model. Curve fits are shown in red. The bivalent analyte model determines kinetic parameters for the first binding event (*K*_D1_, *k*_a1_ and *k*_d1_), representing affinity of binding and for avid binding of the second antibody arm (*K*_D2_, *k*_a2_, and *k*_d2_). B) BLI sensorgrams for binding of low-affinity antibodies to immobilized SARS-CoV-2 RBD-SD1. Data were single reference subtracted and are shown in black. C) BLI sensorgrams showing antibody specificity for SARS-CoV-2 RBD. Binding was measured for antibodies to immobilized SARS-CoV-2 RBD-SD1 or prefusion-stabilized RSV F (DS-Cav1).

**Fig. S5.**
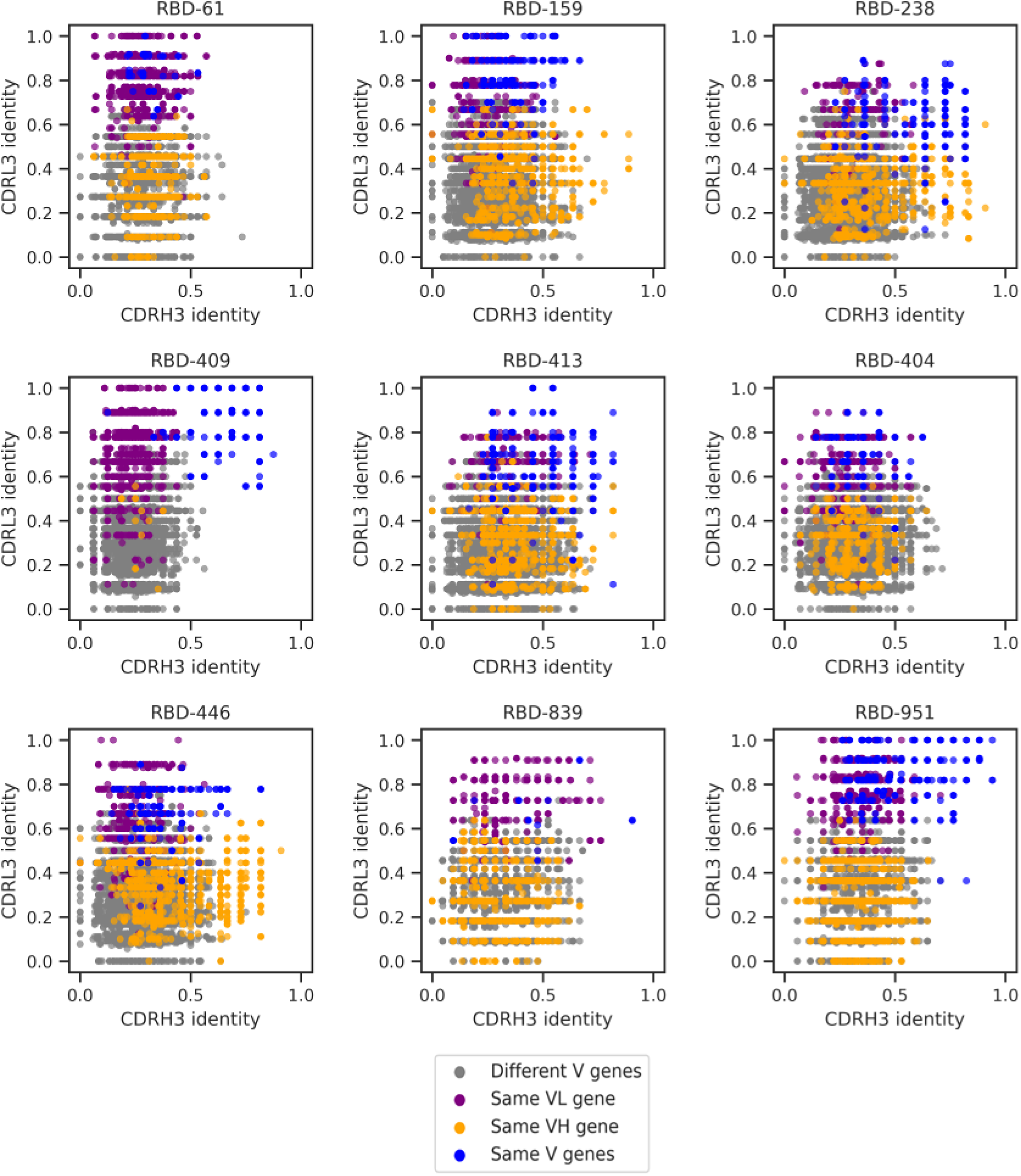
Similarity of designed RBD binders to published RBD-specific antibodies from the CoV-AbDab. Identity is calculated as Levenshtein distance, divided by length of the longer CDR sequence. Pairs are colored based on matching of the V genes.

**Fig. S6.**
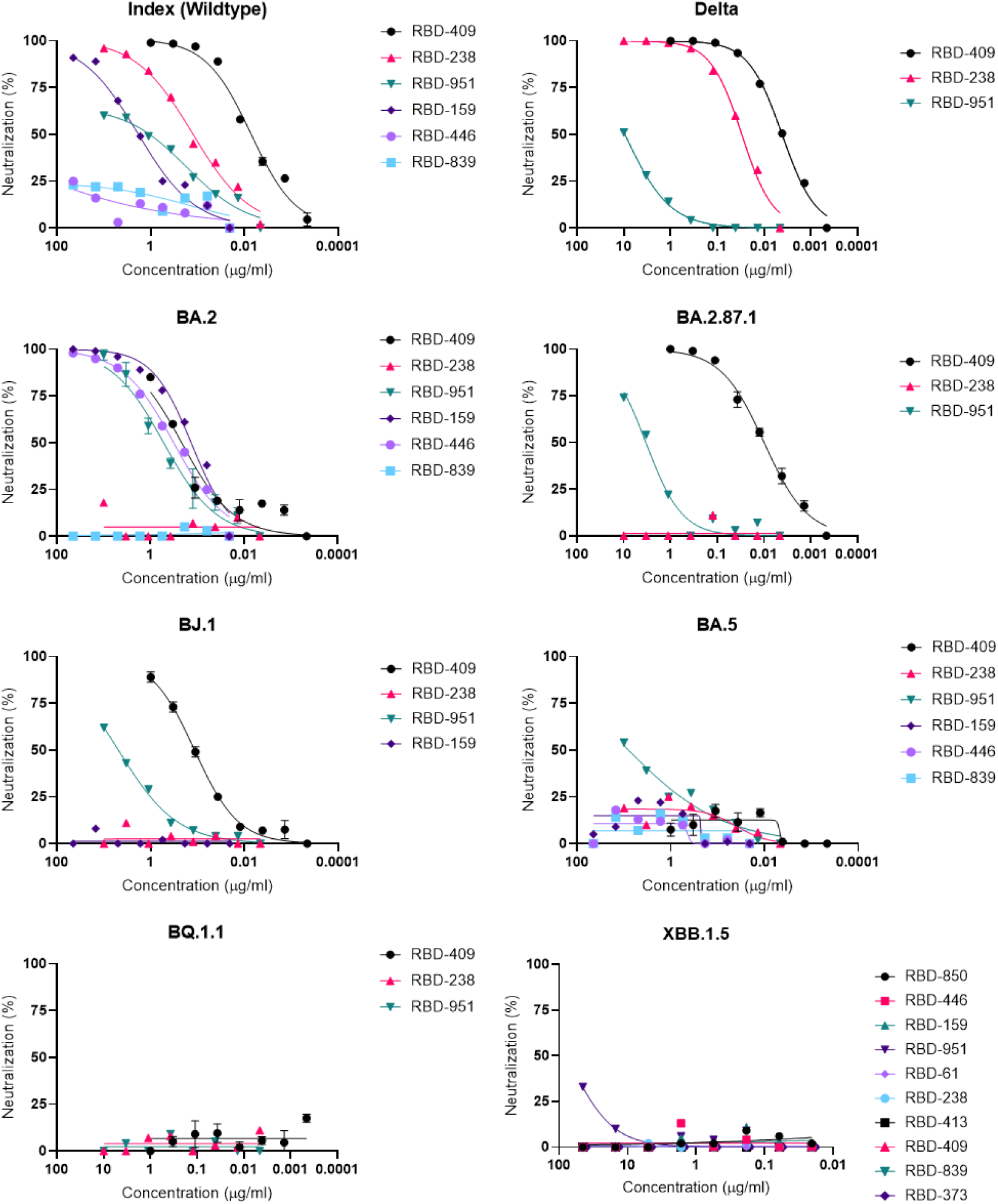
Neutralization dilution curves for RBD-binders against full panel of spike variants.

**Fig. S7.**
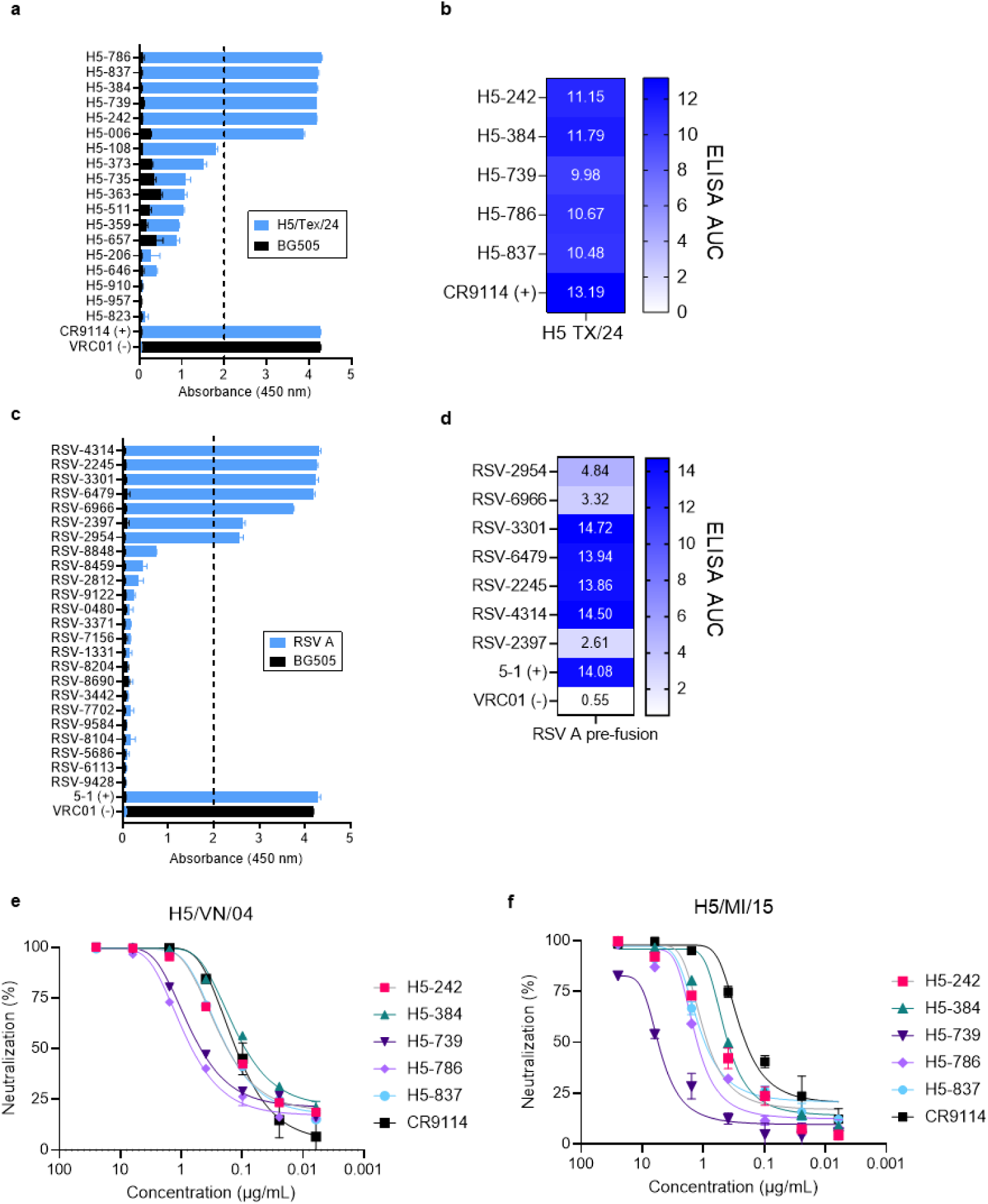
A) Initial ELISA screening for MAGE-generated antibodies against H5/TX/24 hemagglutinin was performed at a concentration of 10μg/mL. Dotted line represents the threshold for further validation. B) ELISA area-under-the-curve (AUC) for H5 prefusion binding antibodies. Calculated from curve shown in Figure 5a. C) Initial ELISA screening for MAGE-generated antibodies against RSV-A prefusion was performed at a concentration of 10μg/mL. Dotted line represents the threshold for further validation. D) ELISA area-under-the-curve (AUC) for RSV-A prefusion binding antibodies. Calculated from curve shown in Figure 6a. Neutralization dilution curves against E) H5/VN/04 and F) H5/MI/15 hemagglutinin.

**Fig. S8.**
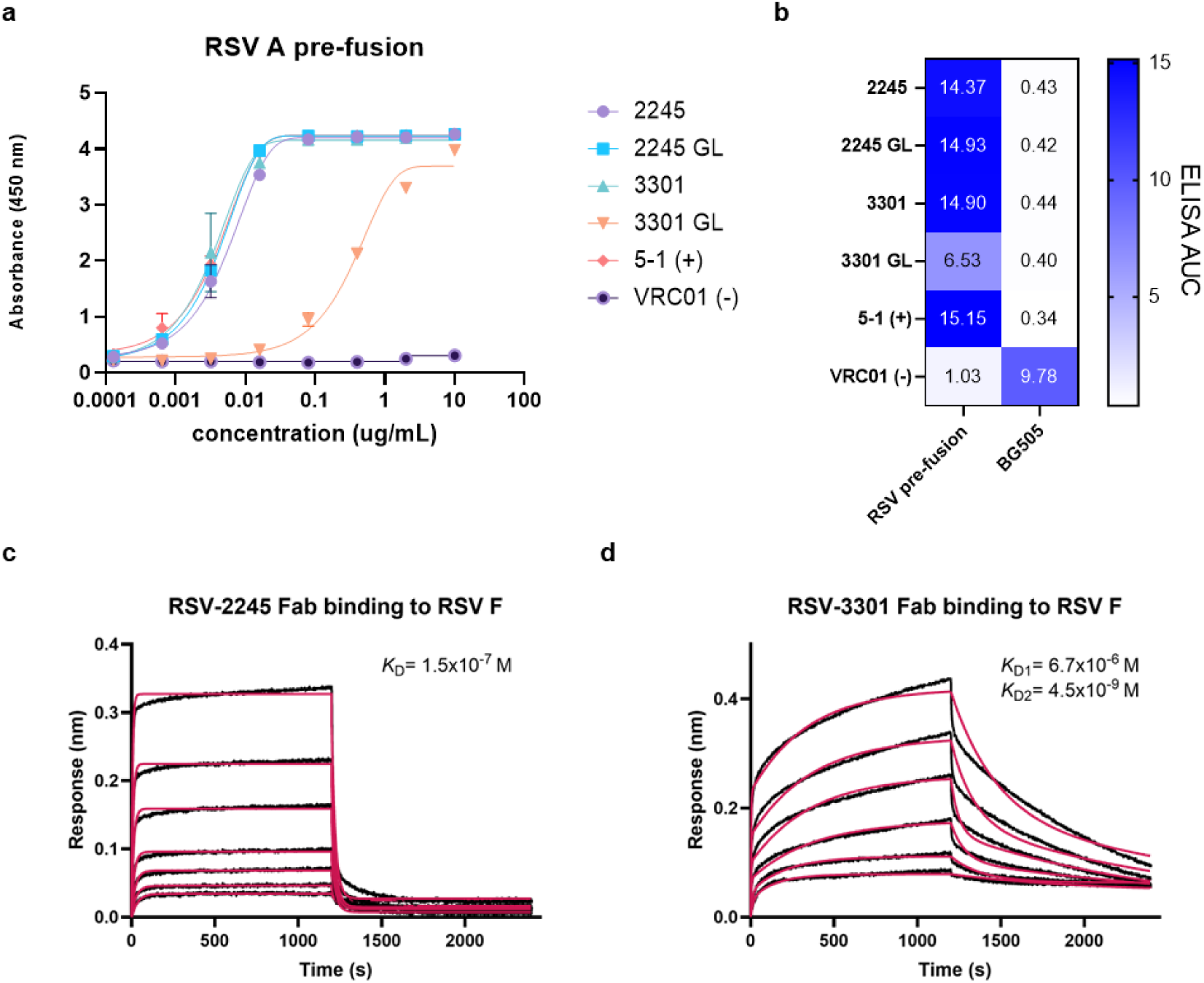
A) RSV-2245 and RSV-3301 were aligned to human germlines, then VH sequences up to CDRH3 were replaced with germline residues and tested for binding by ELISA. B) Area under the curve (AUC) values for germline-reverted ELISA dilution curves. BLI sensorgrams for binding of RSV-2245 Fab (C) and RSV-3301 Fab (D) to immobilized RSV-A F. Data (black) for RSV-2245 binding were fit to a 1:1 binding model to determine binding affinity *(K*_D_). Due to suspected heterogeneity in the epitope targeted by RSV-3301, these data were fit to a heterogeneous ligand model to determine two *K*_D_ values (*K*_D*1*_ and *K*_D*2*_). Curve fits are shown in red.

**Fig. S9.**
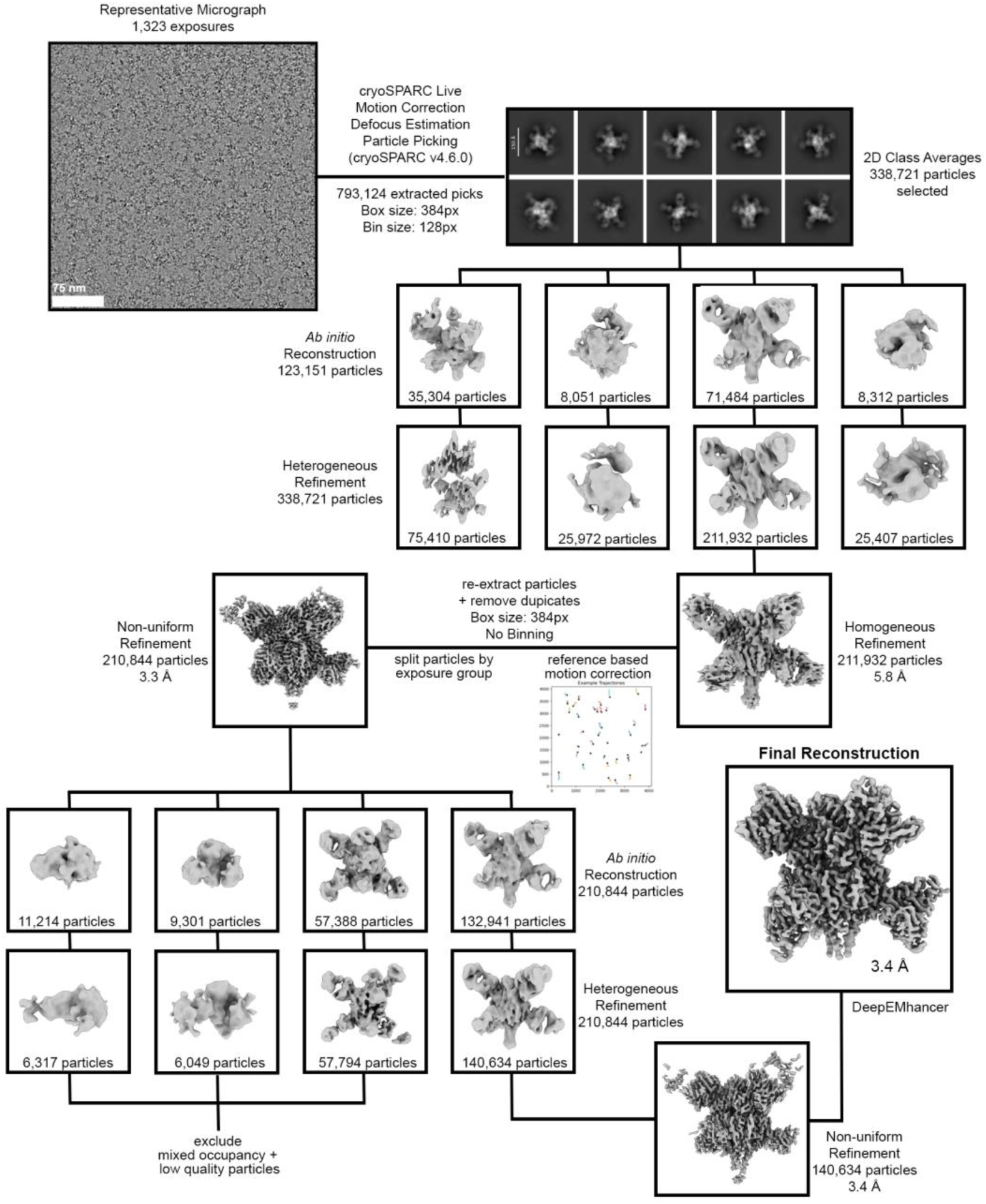
Cryo-EM data processing workflow for RSV prefusion F bound to RSV-2245 and RSV-3301 Fabs.

**Fig. S10.**
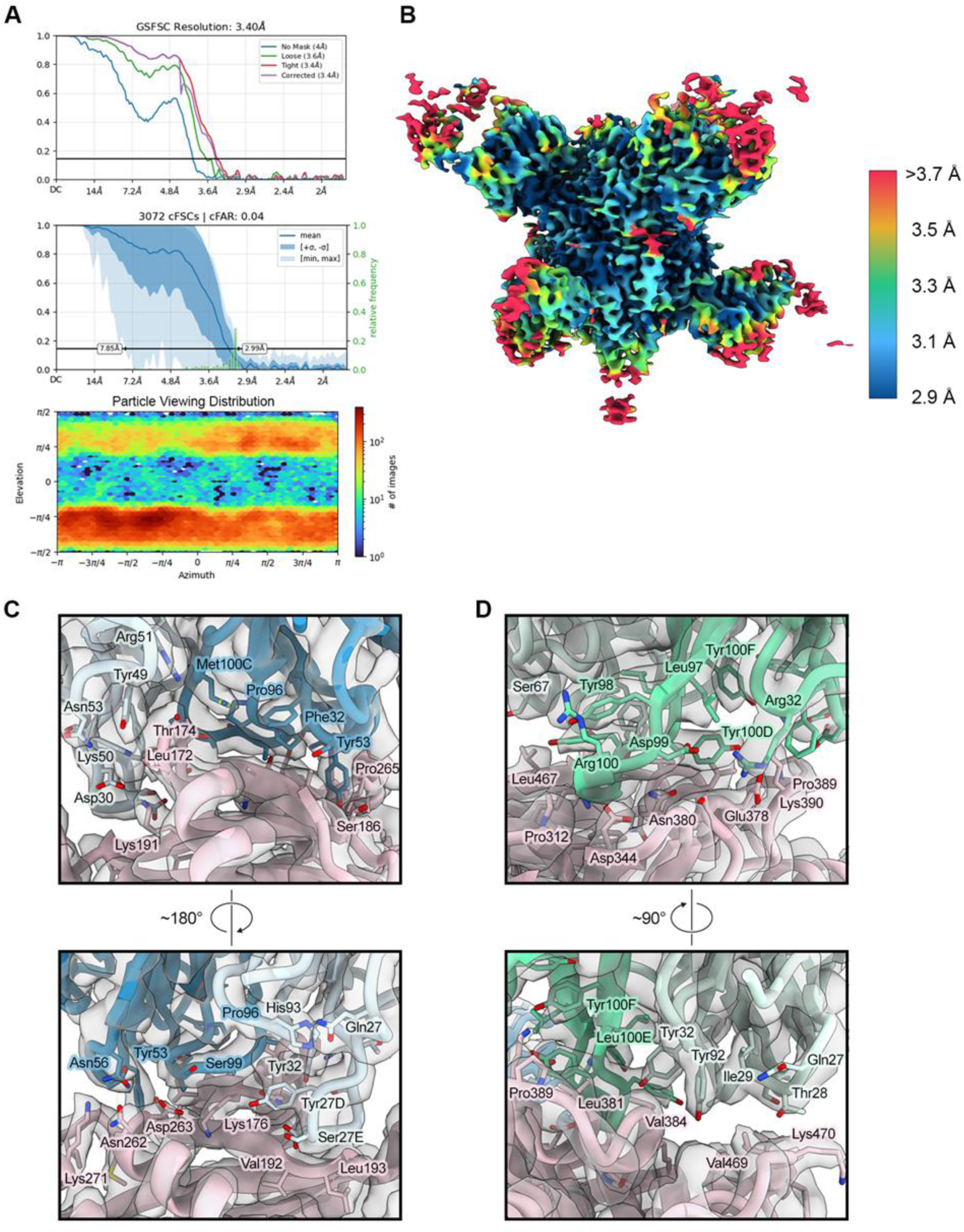
a. Gold-standard Fourier shell correlation, conical Fourier shell correlation, and viewing distribution plots for the RSV F + Fab RSV-2245 + Fab RSV-3301 refinement. b. The final map for the RSV F + Fab RSV-2245 + Fab RSV-3301 complex, colored according to local resolution. c. The binding interface for RSV-2245 and F. The model is shown with F colored pink, the 2245 heavy chain colored blue, and the 2245 light chain colored light blue. The map is partially transparent gray. d. The binding interface for RSV-3301 and F. The 3301 heavy chain is colored green, and the 3301 light chain is colored light green.

**Fig. S11.**
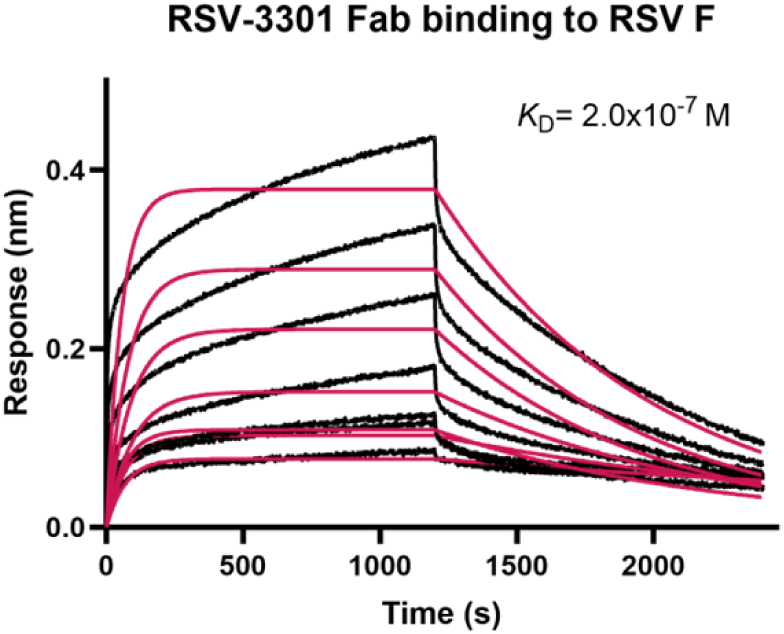
BLI sensorgrams for binding of RSV-3301 Fab to immobilized RSV-A F. Data (black) fit poorly to a 1:1 binding model, leading to an unreliable calculated *K*_D_. Curve fits are shown in red. We performed BLI experiments to measure binding kinetics for RSV-2245 and RSV-3301 Fab binding to immobilized, RSV F trimer (prefusion-stabilized, DS-Cav1) (figure). For RSV-2254, the resulting binding curves displayed rapid saturation during the association phase, followed by a similarly rapid dissociation. A KD of 150 nM was determined by fitting the curves to a 1:1 binding model. RSV-3301 binding resulted in curves demonstrating a fast initial association rate that slowed but did not reach saturation during the 1200-second association step. These curves fit poorly to a 1:1 binding model, shown here. The RSV F trimer has been shown to transiently open, which can affect the accessibility of some epitopes. Additionally, DS-Cav1 maintains some prefusion instability that may lead to pre- and postfusion conformations of F immobilized on the sensortip. Because the RSV-3301 epitope is largely conserved in postfusion F, simultaneous binding of Fab to pre- and postfusion trimers might be observed. These considerations led us to fit the curves to a heterogeneous ligand model, resulting in apparent KD values (KD1 and KD2) of 6.7 μM and 4.5 nM, respectively.

**Table S3.**
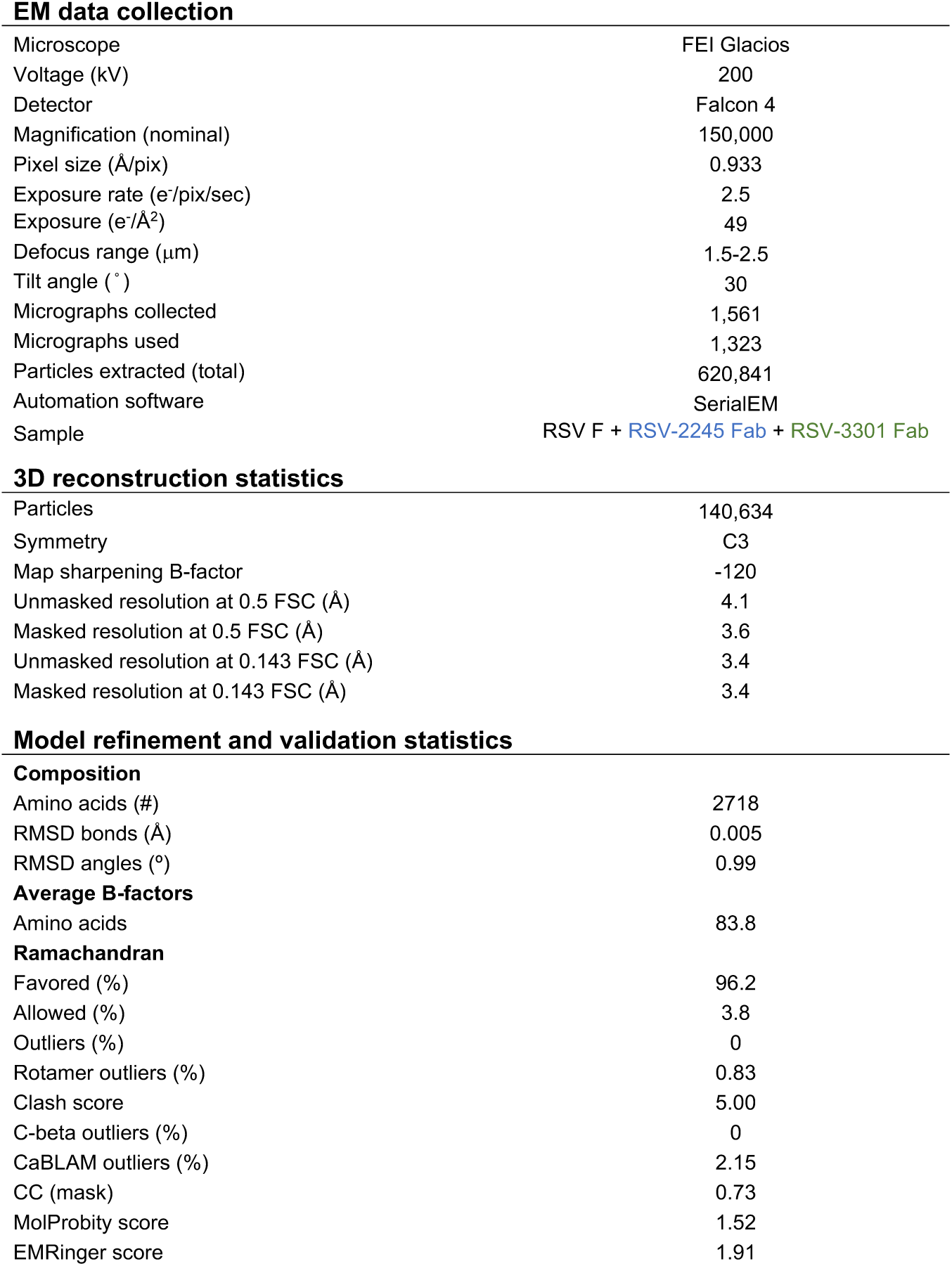
Cryo-EM data collection, processing, and PDB model validation.

**EM data collection**
|  |  |
| --- | --- |
| Microscope | FEI Glacios |
| Voltage (kV) | 200 |
| Detector | Falcon 4 |
| Magnification (nominal) | 150,000 |
| Pixel size (Å/pix) | 0.933 |
| Exposure rate (e<sup>-</sup>/pix/sec) | 2.5 |
| Exposure (e<sup>-</sup>/Å<sup>2</sup>) | 49 |
| Defocus range (µm) | 1.5-2.5 |
| Tilt angle (°) | 30 |
| Micrographs collected | 1,561 |
| Micrographs used | 1,323 |
| Particles extracted (total) | 620,841 |
| Automation software | SerialEM |
| Sample | RSV F + RSV-2245 Fab + RSV-3301 Fab |
| **3D reconstruction statistics** |  |
| Particles | 140,634 |
| Symmetry | C3 |
| Map sharpening B-factor | -120 |
| Unmasked resolution at 0.5 FSC (Å) | 4.1 |
| Masked resolution at 0.5 FSC (Å) | 3.6 |
| Unmasked resolution at 0.143 FSC (Å) | 3.4 |
| Masked resolution at 0.143 FSC (Å) | 3.4 |
| **Model refinement and validation statistics****Composition** |  |
| Amino acids (#) | 2718 |
| RMSD bonds (Å) | 0.005 |
| RMSD angles (°) | 0.99 |
| **Average B-factors** |  |
| Amino acids | 83.8 |
| Favored (%) | 96.2 |
| Allowed (%) | 3.8 |
| Outliers (%) | 0 |
| Rotamer outliers (%) | 0.83 |
| Clash score | 5.00 |
| C-beta outliers (%) | 0 |
| CaBLAM outliers (%) | 2.15 |
| CC (mask) | 0.73 |
| MolProbity score | 1.52 |
| EMRinger score | 1.91 |

## Notes

### Summary of Updates

Updating funding information, fixed typo in Figure 5 caption.

