## Supplemental figures for "Generation of antigen-specific paired chain antibody sequences using large language models"

### Slide 1
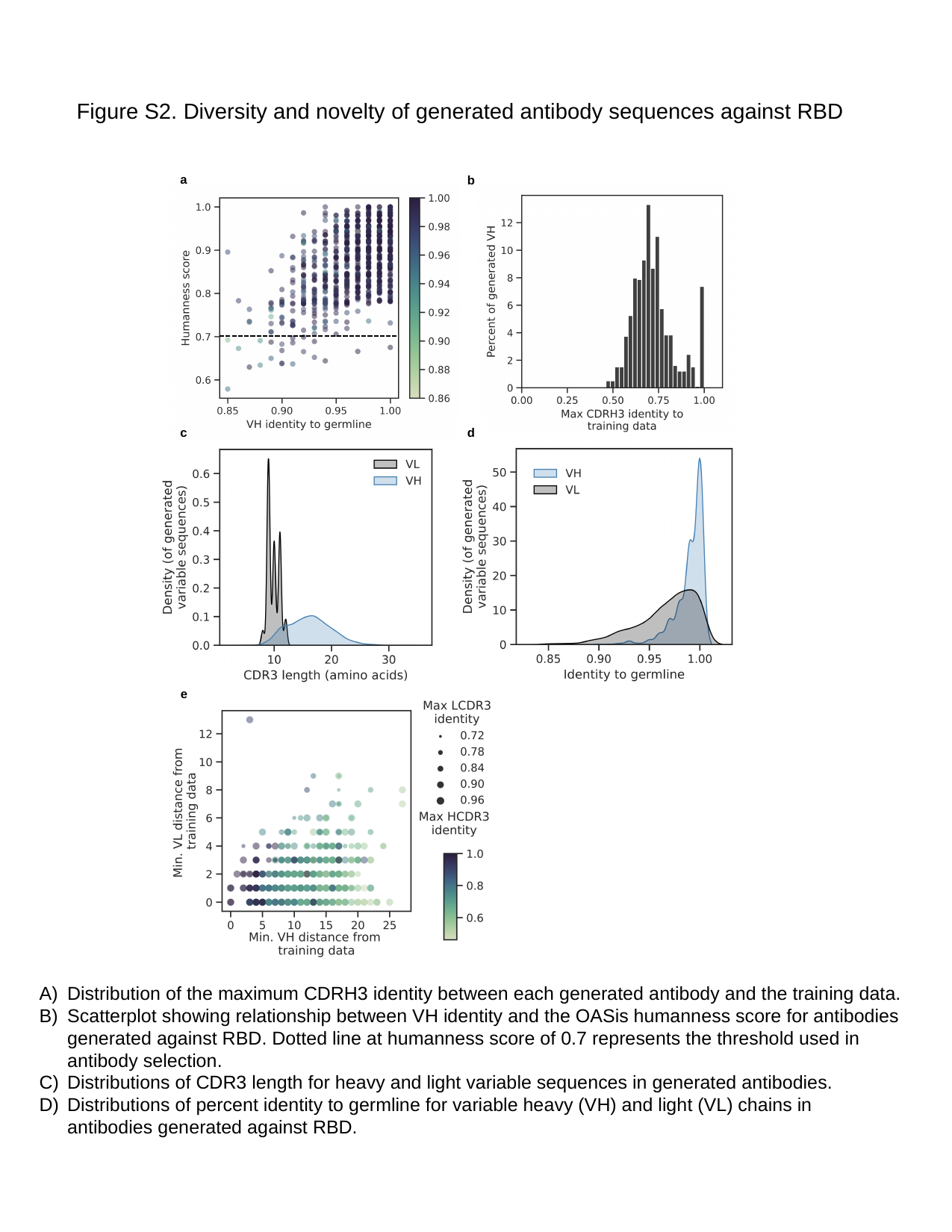

#### Figure S2. Diversity and novelty of generated antibody sequences against RBD
a
b
c
d
e
Distribution of the maximum CDRH3 identity between each generated antibody and the training data.
Scatterplot showing relationship between VH identity and the OASis humanness score for antibodies generated against RBD. Dotted line at humanness score of 0.7 represents the threshold used in antibody selection.
Distributions of CDR3 length for heavy and light variable sequences in generated antibodies.
Distributions of percent identity to germline for variable heavy (VH) and light (VL) chains in antibodies generated against RBD.
E) Based on the comparison in C, the distance to the closest training example for the generated VH and VL sequences is shown. Size of the points represents the maximum LCDR3 to any training sequence, and color represents the maximum HCDR3 to any training sequence.

### Slide 2
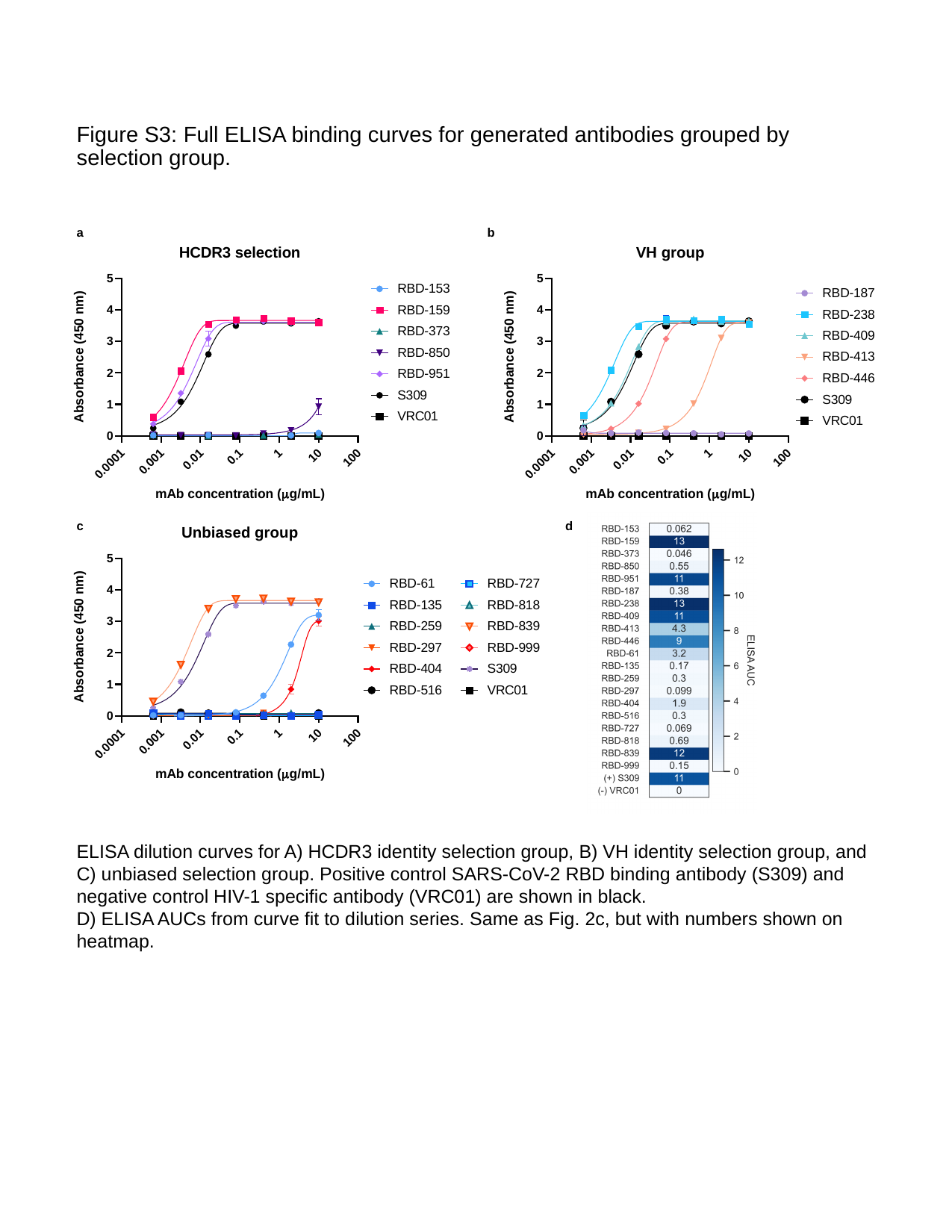

Figure S3: Full ELISA binding curves for generated antibodies grouped by selection group.
a
b
c
d
ELISA dilution curves for A) HCDR3 identity selection group, B) VH identity selection group, and C) unbiased selection group. Positive control SARS-CoV-2 RBD binding antibody (S309) and negative control HIV-1 specific antibody (VRC01) are shown in black.
D) ELISA AUCs from curve fit to dilution series. Same as Fig. 2c, but with numbers shown on heatmap.

### Slide 3
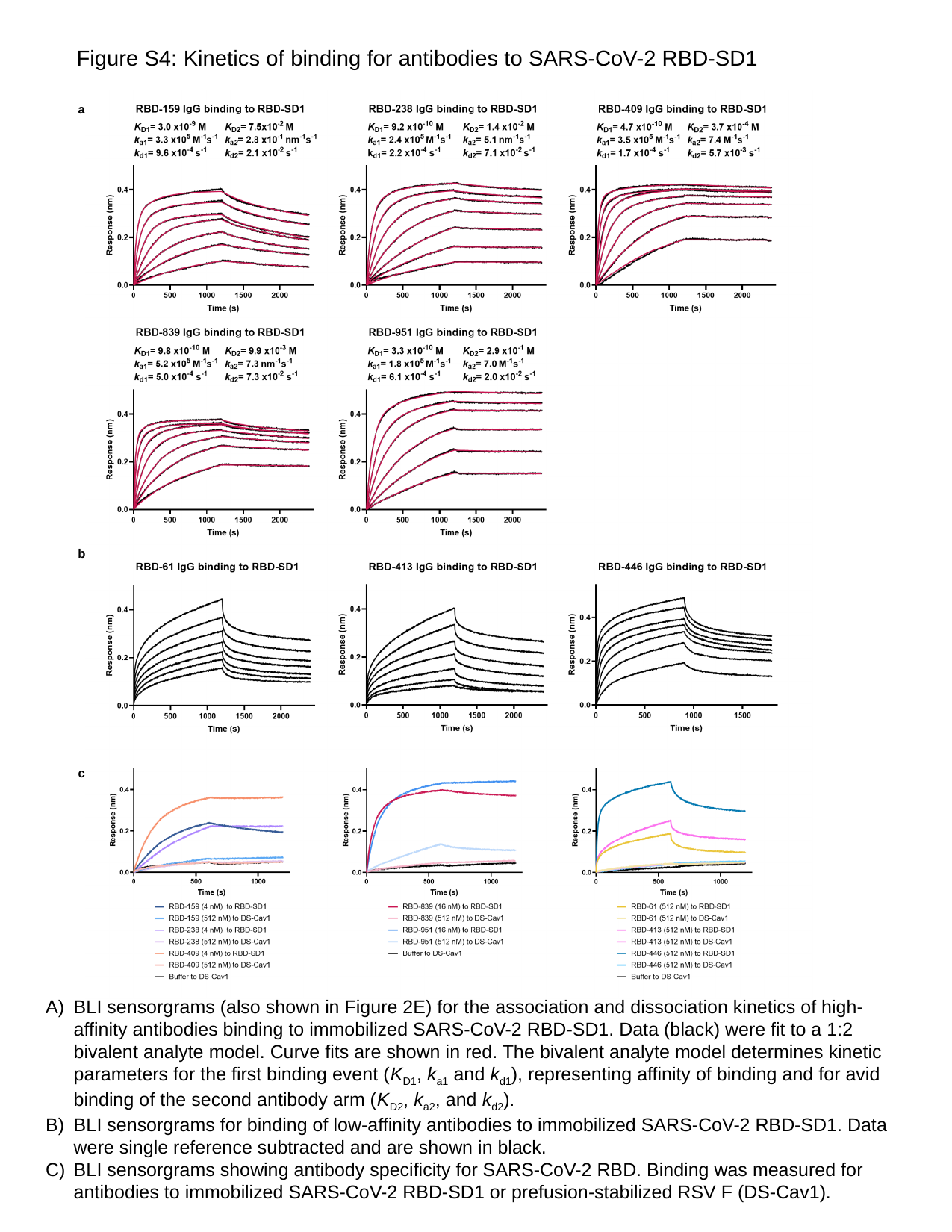

Figure S4: Kinetics of binding for antibodies to SARS-CoV-2 RBD-SD1
a
b
c
BLI sensorgrams (also shown in Figure 2E) for the association and dissociation kinetics of high-affinity antibodies binding to immobilized SARS-CoV-2 RBD-SD1. Data (black) were fit to a 1:2 bivalent analyte model. Curve fits are shown in red. The bivalent analyte model determines kinetic parameters for the first binding event (K_D1_, k_a1_ and k_d1_), representing affinity of binding and for avid binding of the second antibody arm (K_D2_, k_a2_, and k_d2_).
BLI sensorgrams for binding of low-affinity antibodies to immobilized SARS-CoV-2 RBD-SD1. Data were single reference subtracted and are shown in black.
BLI sensorgrams showing antibody specificity for SARS-CoV-2 RBD. Binding was measured for antibodies to immobilized SARS-CoV-2 RBD-SD1 or prefusion-stabilized RSV F (DS-Cav1).

### Slide 4
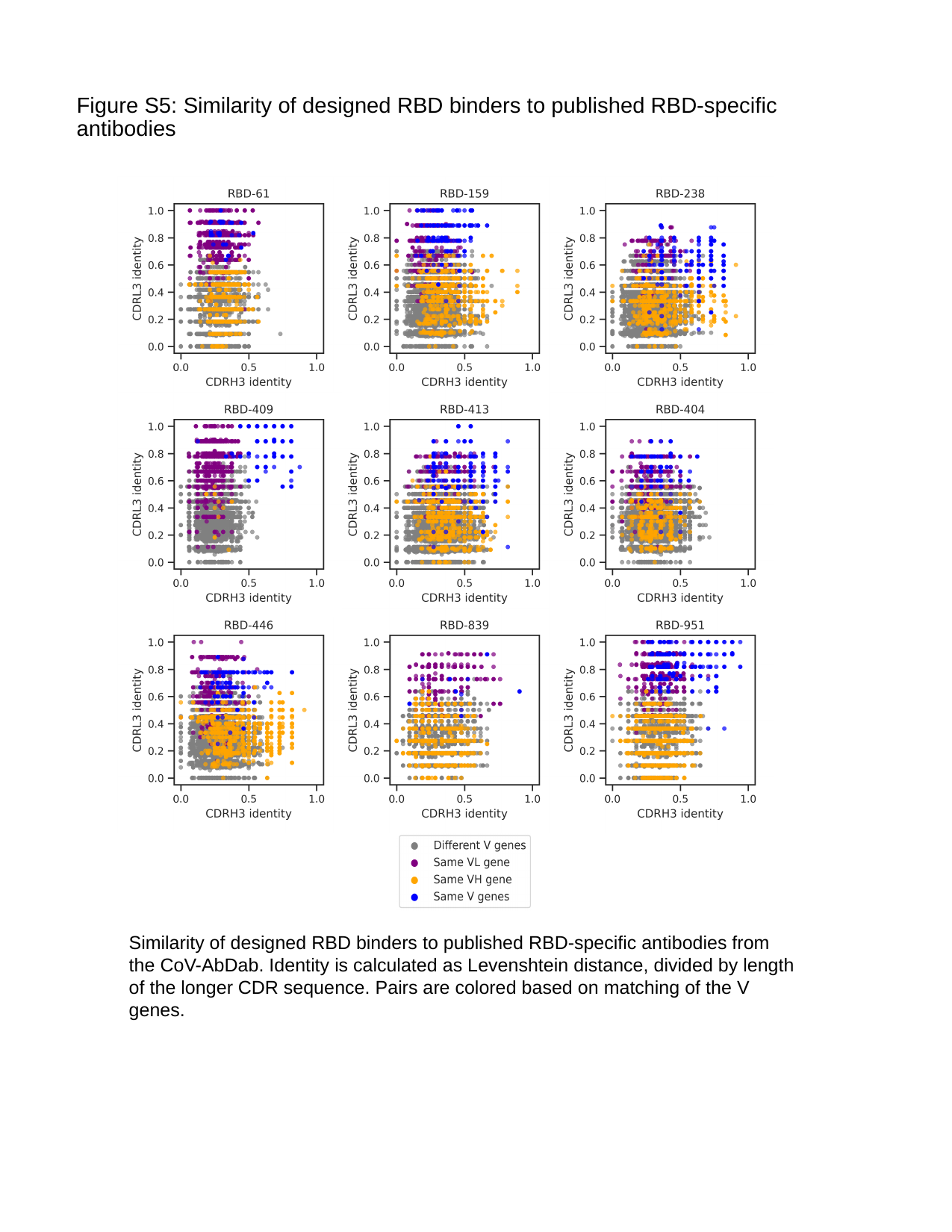

Figure S5: Similarity of designed RBD binders to published RBD-specific antibodies
Similarity of designed RBD binders to published RBD-specific antibodies from the CoV-AbDab. Identity is calculated as Levenshtein distance, divided by length of the longer CDR sequence. Pairs are colored based on matching of the V genes.

### Slide 5
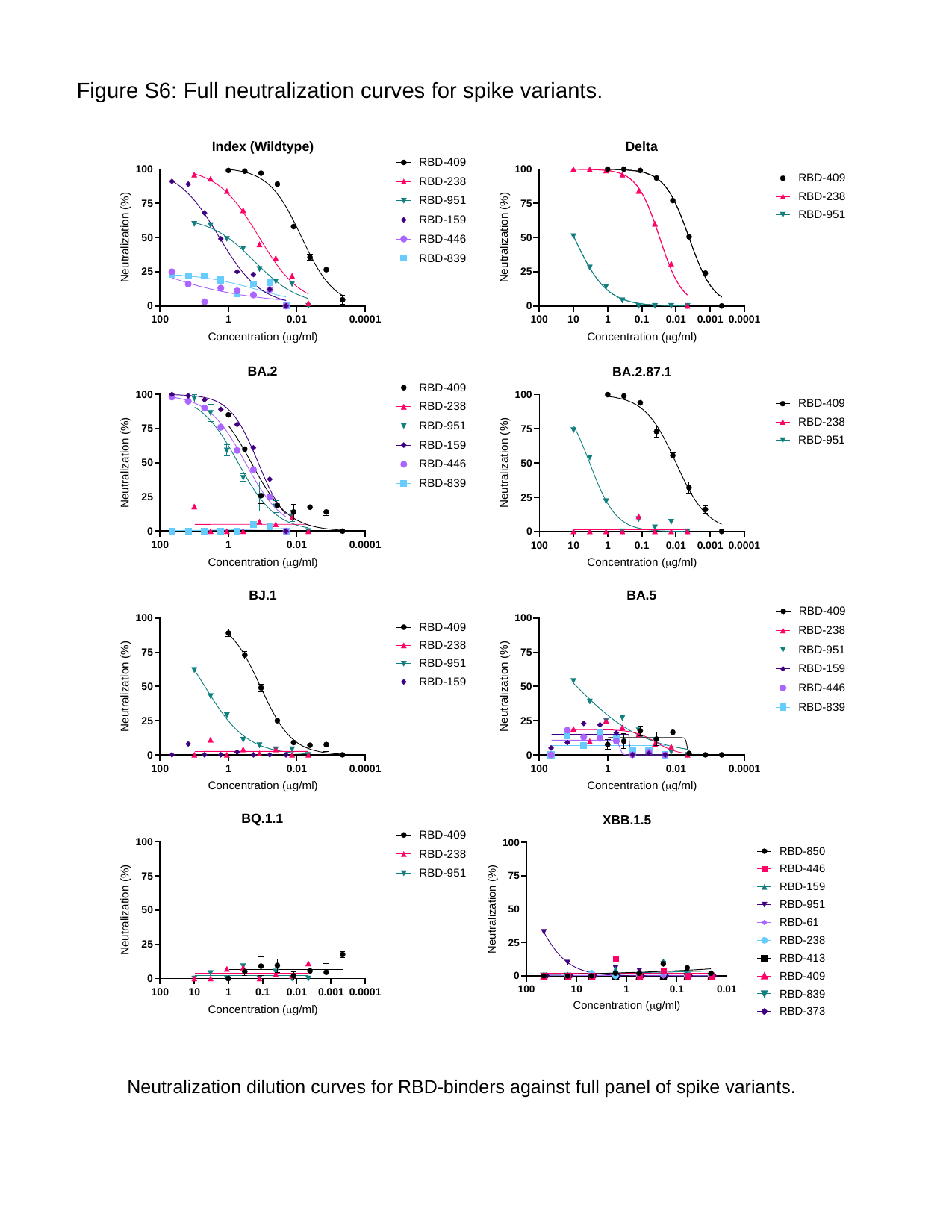

Figure S6: Full neutralization curves for spike variants.
Neutralization dilution curves for RBD-binders against full panel of spike variants.

### Slide 6
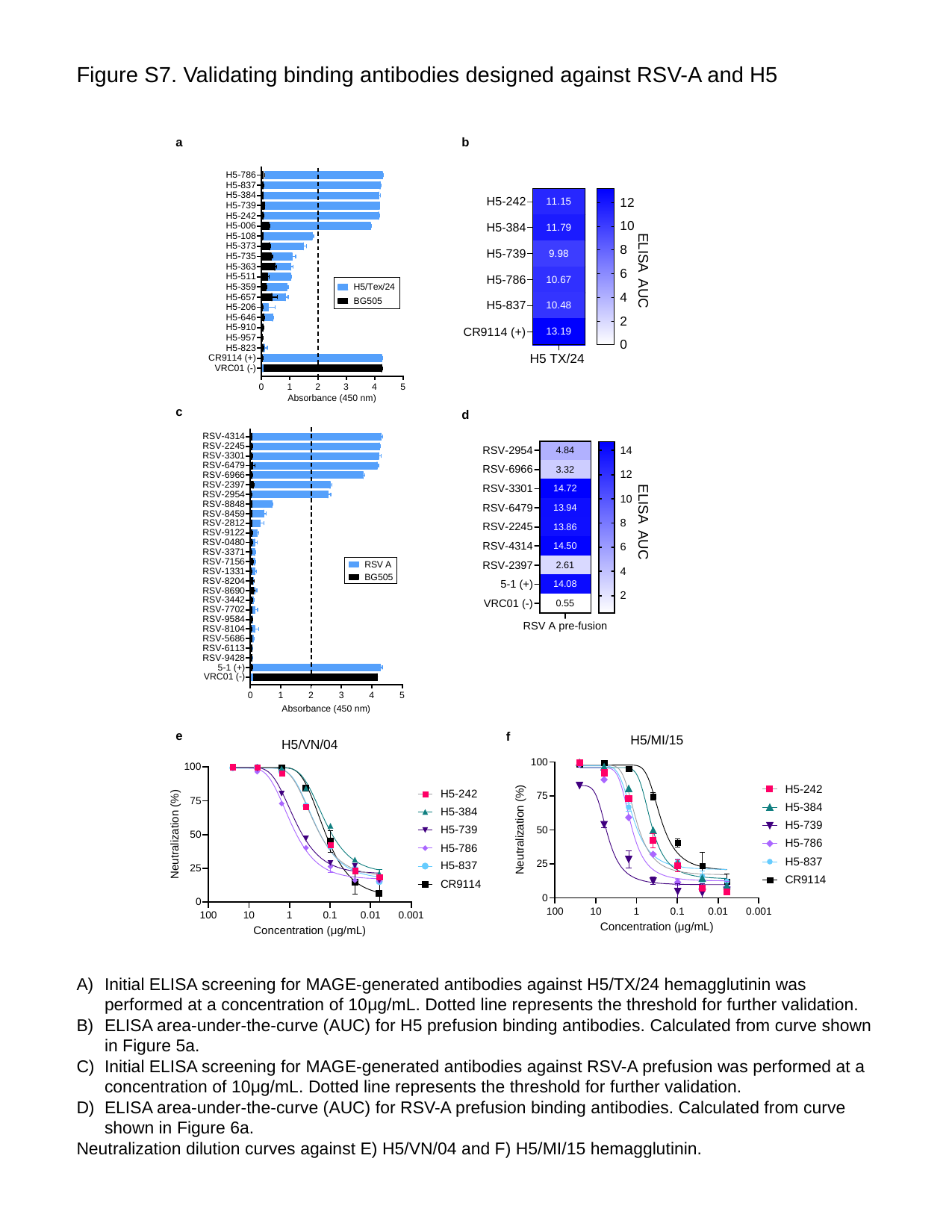

Figure S7. Validating binding antibodies designed against RSV-A and H5
a
b
ELISA AUC
c
d
ELISA AUC
e
f
Initial ELISA screening for MAGE-generated antibodies against H5/TX/24 hemagglutinin was performed at a concentration of 10μg/mL. Dotted line represents the threshold for further validation.
ELISA area-under-the-curve (AUC) for H5 prefusion binding antibodies. Calculated from curve shown in Figure 5a.
Initial ELISA screening for MAGE-generated antibodies against RSV-A prefusion was performed at a concentration of 10μg/mL. Dotted line represents the threshold for further validation.
ELISA area-under-the-curve (AUC) for RSV-A prefusion binding antibodies. Calculated from curve shown in Figure 6a.
Neutralization dilution curves against E) H5/VN/04 and F) H5/MI/15 hemagglutinin.

### Slide 7
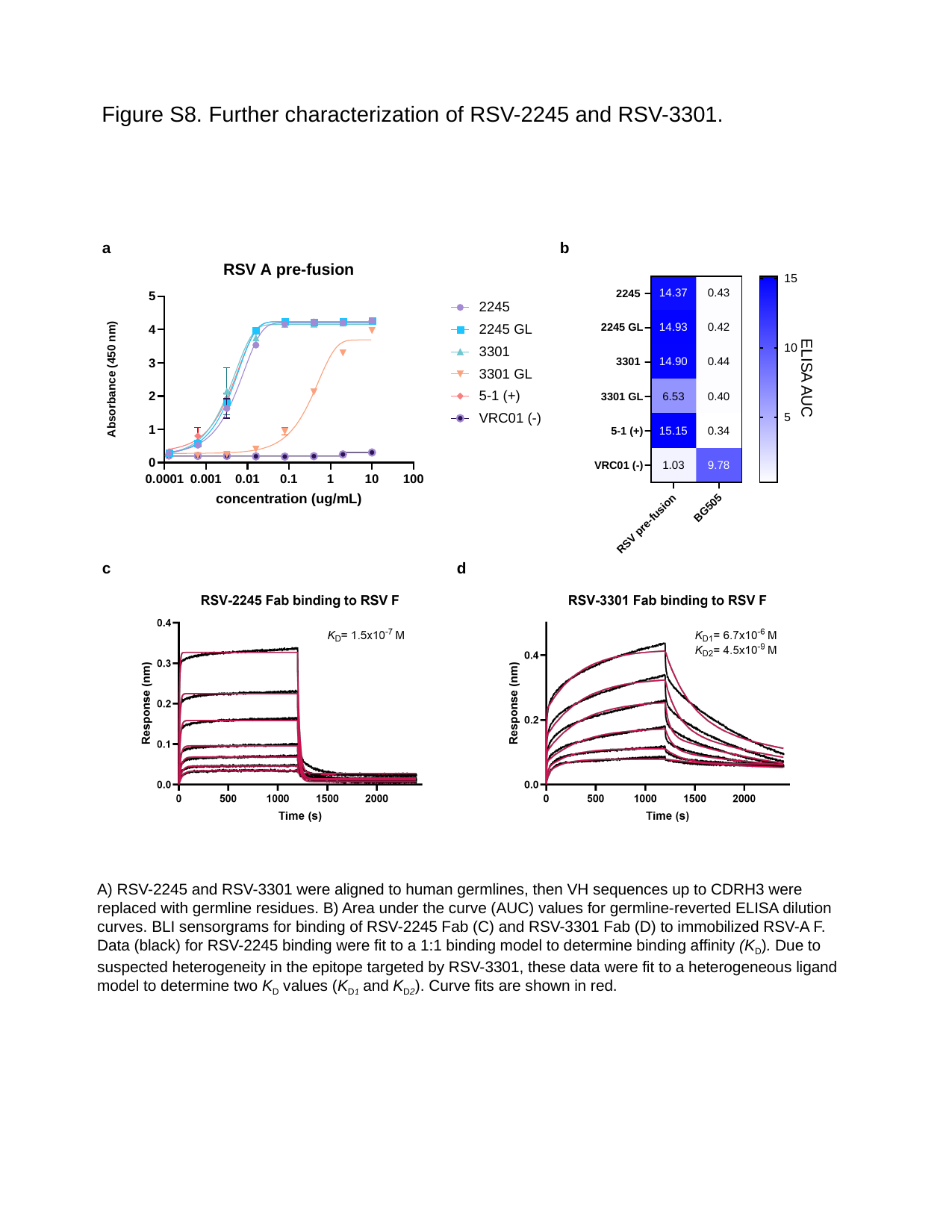

Figure S8. Further characterization of RSV-2245 and RSV-3301.
a
b
ELISA AUC
c
d
A) RSV-2245 and RSV-3301 were aligned to human germlines, then VH sequences up to CDRH3 were replaced with germline residues. B) Area under the curve (AUC) values for germline-reverted ELISA dilution curves. BLI sensorgrams for binding of RSV-2245 Fab (C) and RSV-3301 Fab (D) to immobilized RSV-A F. Data (black) for RSV-2245 binding were fit to a 1:1 binding model to determine binding affinity (K_D_). Due to suspected heterogeneity in the epitope targeted by RSV-3301, these data were fit to a heterogeneous ligand model to determine two K_D_ values (K_D1_ and K_D2_). Curve fits are shown in red.
